# Age-specific regulation of sociability by hypothalamic Agrp neurons

**DOI:** 10.1101/2025.05.05.652061

**Authors:** Onur Iyilikci, Lucas Kim, Marcelo R. Zimmer, Jeremy Bober, Yuexuan Li, Macy Pelts, Gustavo M. Santana, Marcelo O. Dietrich

**Affiliations:** Laboratory of Physiology of Behavior, Department of Comparative Medicine, Yale School of Medicine, New Haven, CT, USA; Yale Center for Molecular and Systems Metabolism, Yale School of Medicine, New Haven, CT, USA; Department of Neuroscience, Yale School of Medicine, New Haven, CT, USA; University of Ozarks, Clarksville, AR, USA

## Abstract

Social isolation enhances sociability, suggesting that social behavior is maintained through a homeostatic mechanism. Further, mammalian social needs shift dramatically from infancy through adolescence into adulthood, raising the question of whether the neural mechanisms governing this homeostatic regulation evolve across developmental stages. Here, we show that agouti-related peptide (Agrp) neurons, which regulate hunger in adults, are activated by social isolation from weaning through adolescence but not in adulthood. Importantly, the activity of these neurons is critical for social behavior during adolescence: inhibiting Agrp neurons reduced isolation-induced sociability in juveniles but not in adults, and Agrp neuron activation promoted sociability only in young mice. After isolation, reunion with siblings or other conspecifics, but not unfamiliar adult males, rapidly decreased neuronal activity in juveniles, an effect requiring intact olfaction. These findings identify Agrp neurons as a key component of the circuitry governing age-specific social homeostasis.

## Introduction

Humans and other mammals exhibit heightened sociability following periods of social isolation, suggesting that social behavior is regulated by a homeostatic mechanism that monitors and restores social needs. While this concept has intuitive appeal and has been widely used to explain sociability^1-10^, the neural substrates underlying ‘social homeostasis’ remain poorly understood. Adding complexity, in mammals, social needs change drastically across different periods of life, as individuals transition from infancy into adolescence and ultimately to adulthood. For instance, infants’ social relationships shift from infant-mother interactions^11^ to juvenile play^10^ and eventually to adult social behaviors. Despite the critical nature of these developmental transitions, how the brain regulates social needs at different ages remains elusive. Agouti-related peptide-expressing (Agrp) neurons in the hypothalamus are best known for their role in hunger regulation in adults. Nutrient deprivation increases their firing rate^12-15^, while food ingestion reduces their activity^16-19^, effectively making Agrp neuron activity a readout of an animal’s overall need for food^20^. Interestingly, in neonatal mice, Agrp neurons behave differently^21-23^. For example, separation of ten-days-old mouse pups from the nest activates these neurons, even when the pups are fed^23^. Separation in the presence of a non-lactating female blunts this activation, suggesting that Agrp neurons in neonates do not solely respond to the nutritional state of the pup^24^. Based on these observations, we reasoned that Agrp neurons may function as key regulators of age-specific social needs. Here, we discovered that Agrp neurons specifically regulate social homeostasis during a sensitive period spanning from infancy through adolescence. Their involvement in social homeostasis does not extend into adulthood, revealing a transient, age-specific role for this canonical feeding circuit in social behavior of mice.

## Results

### Social isolation activates Agrp neurons in juveniles

We first set out to test whether social isolation activates Agrp neurons in juvenile mice. In adolescence (see **Fig 1A** for developmental periods in mice^25,26^), social isolation strongly increases rebound sociability^2,27-29^ and social interactions are typically non-sexual and non-aggressive, reflecting the unique social demands of juveniles^25,27^.

**Figure 1:**
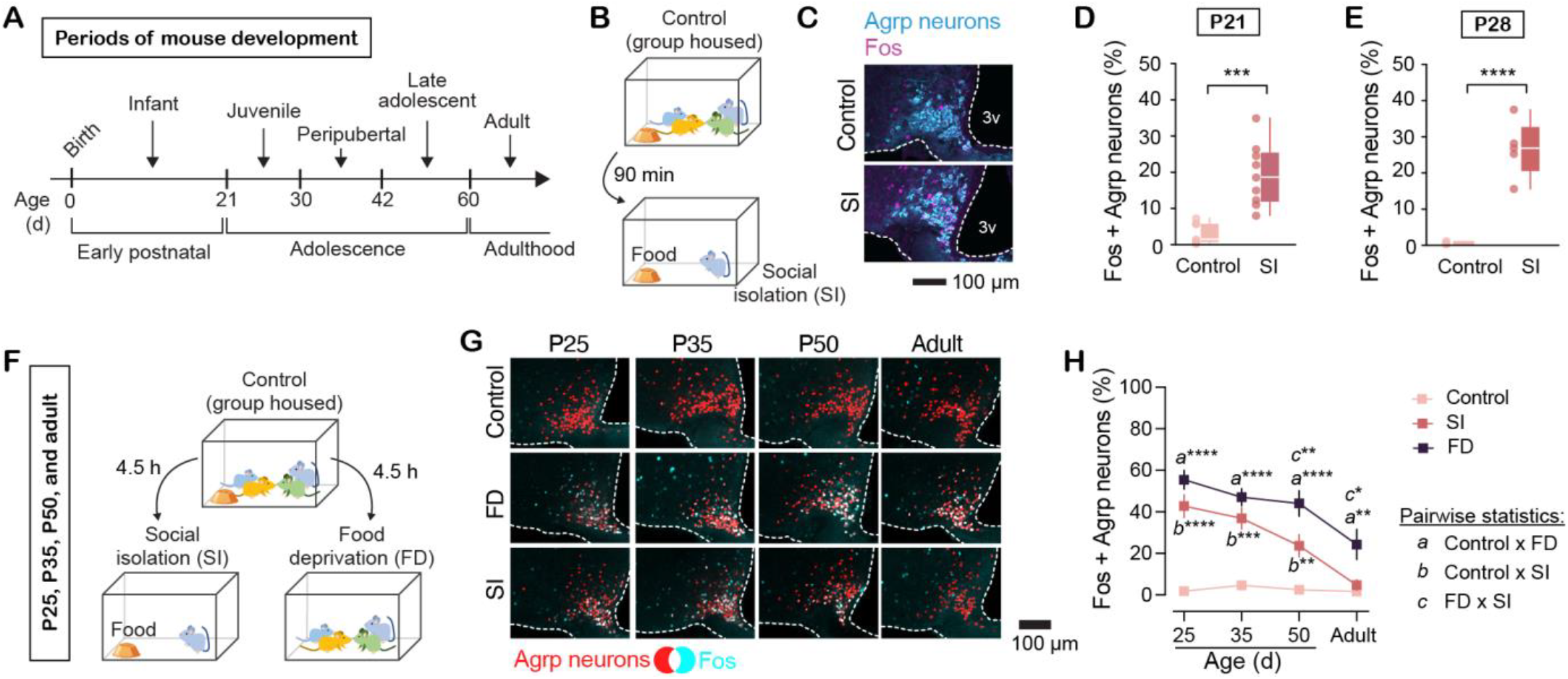
Social isolation activates Agrp neurons during adolescence. **(A)** Periods of mouse development and nomenclature used in this manuscript. **(B)** Juvenile mice (P21 and P28) were socially isolated (SI) for 90 minutes with ad libitum food. **(C)** Representative images of Fos labeling in Agrp neurons of control and SI juvenile Agrp^HA^ mice (see Methods). **(D)** Agrp neurons positive for Fos at P21 (nest, n = 7; isolation, n = 9; *t*_14_ = 4.69, *P*_2-tailed_ = 0.0003) and **(E)** P28 (nest, n = 5; isolation, n = 5; *t*_8_ = 7.45, *P*_2-tailed_ = 0.00007). **(F)** Protocol to compare the activation of Agrp neurons in response to social isolation with ad libitum food (SI) and food deprivation in social group (FD) across ages: P25, P35, P50, and adult, P65+; mice were SI or FD for 4.5 hours. **(G)** Representative images of Fos labeling in Agrp neurons of control, SI, and FD mice at different ages (tdTomato labels the nuclei of Agrp neurons). **(H)** Agrp neurons positive for Fos in control, SI, and FD mice at different ages [ccontrol: P25, n = 4; P35, n = 4; P50, n = 4; and adult, n = 6; SI: P25, n = 4; P35, n = 4; P50, n = 7; and adult, n = 5; FD: P25, n = 4; P35, n = 3; P50, n = 5; and adult, n = 5]; effect of age: *F*_3, 43_ = 14.40, *P* = 0. 000001; group: *F*_2, 43_ = 68.93, *P* < 10^-13^; interaction: *F*_6, 43_ = 3.68, *P* = 0. 004. In D-E: statistical analyses using unpaired *t* test. In H, statistical analyses using two-way ANOVA with age and group as factors followed by Holm-Sidak’s multiple comparisons test (indicated by the letters *a, b, c* for each pairwise comparison). Symbols represent individual data. Box plot denotes minimum, first quartile, median, third quartile, and maximum values. In H, symbols represent mean ± sem. * *P* < 0.05; ** *P* < 0.01; *** *P* < 0.001; **** *P* < 0.0001.

We measured activation of Agrp neurons in ad libitum fed mice after social isolation (90 minutes) using Fos. We found that both at P21 (at weaning) and P28 (a week after weaning), social isolation significantly activates Agrp neurons (**Fig. 1B-E**). We next compared the effect of social isolation (in the presence of ad libitum food) to the effect of food deprivation (in social group) from adolescence to adulthood (P25, P35, P50, and adults, P60 or older) (**Fig. 1F**). For this experiment, we socially isolated or food deprived mice for 4.5 hours, as this period of food deprivation strongly activates Agrp neurons in adults^30,31^. We found that social isolation significantly activates Agrp neurons during adolescence, but not in adults. The effect in young mice was of similar magnitude as the activation of Agrp neurons by food deprivation in socially housed mice (**Fig. 1G-H**). Together, these results demonstrate that during adolescence, when social behaviors are not necessary to meet basic needs as they do in infancy, social isolation strongly activates Agrp neurons, suggesting a potential role for these neurons in the maintenance of social homeostasis during adolescence.

### Agrp neurons are required for isolation-induced sociability in juveniles

Our Fos experiments suggest that Agrp neurons may signal social needs, similar to how they signal the need for food in adults^20^. If this is true, changing the activity of these neurons should have an effect on social behaviors (**Fig. 2A**).

**Figure 2:**
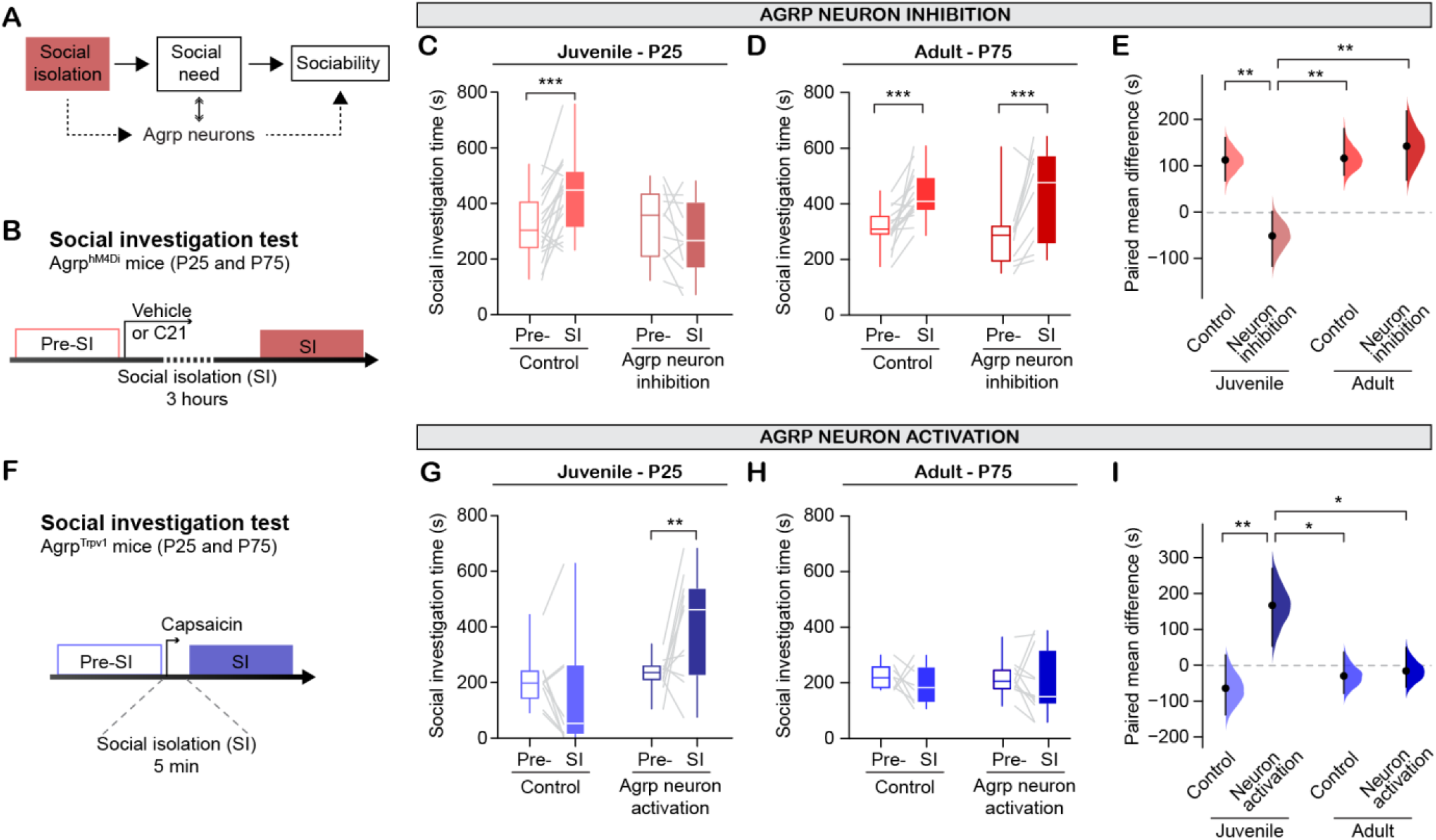
Modulation of social investigation by Agrp neurons in juvenile mice, but not in adults. **(A)** Model by which social isolation (SI) increases social need and sociability; Agrp neurons may encode social need and, consequently, sociability. **(B)** Experimental design to test the requirement of Agrp neuron activity to increase social investigation after social isolation. Mice expressing hM4Di in Agrp neurons were randomly injected with vehicle or C21 at specific ages (P25 or P75) immediately before social isolation (for 3 hours). Mice with no expression of hM4Di (missed viral injection) and that received C21 were used as a control group for the effects of C21 and pooled together in the control group. **(C)** Effect of social isolation (*F*_1, 26_ = 8.11, *P* = 0.008) and neuron inhibition (*F*_2, 26_ = 1.83, *P* = 0.17) on social investigation (interaction, *F*_2, 26_ = 8.00, *P* = 0.002) in juveniles (control, n = 18; Agrp neuron inhibition, n = 11). **(D)** Similar to C, but for adult mice (effect of social isolation: *F*_1, 23_ = 27.38, *P* < 0.0001; neuron inhibition: *F*_2, 23_ = 0.41, *P* = 0.66; interaction: *F*_2, 23_ = 0.16, *P* = 0.84; control, n = 16; Agrp neuron inhibition, n = 10). **(E)** Estimation plot displaying the paired mean difference (± 95% CI) of social investigation time after SI minus pre-SI (effect of age: *F*_1, 51_ = 10.52, *P* = 0. 002; neuron inhibition: *F*_1, 51_ = 5.11, *P* = 0.02; interaction: *F*_1, 51_ = 9.72, *P* = 0. 003). **(F)** Experimental design to test the capacity of Agrp neurons to increase sociability. Mice expressing Trpv1 in Agrp neurons, and their littermate controls that do not express Trpv1, were injected with capsaicin to activate Agrp neurons immediately after a baseline social investigation test and then socially isolated for 5 minutes before repeating the test. **(G)** Effect of social isolation for five minutes (*F*_1, 18_ = 1.75, *P* = 0.2) and neuron activation (*F*_1, 18_ = 6.01, *P* = 0.02) on social investigation (interaction: *F*_1, 18_ = 8.88, *P* = 0.008) in juveniles (control, n = 9; *Agrp*^Trpv1^, n = 11). **(H)** Similar to G but for adult mice (effect of social isolation, *F*_1, 18_ = 1.20, *P* = 0.28; neuron activation: *F*_1, 18_ = 0.006, *P* = 0.93; interaction: *F*_1, 18_ = 0.79, *P* = 0.79; control, n = 8; *Agrp*^Trpv1^, n = 12). **(I)** Estimation plot displaying the paired mean difference (± 95% CI) of social investigation time after SI minus pre-SI (effect of age: *F*_1, 36_ = 2.84, *P* = 0.10; neuron activation: *F*_1, 36_ = 7.35, *P* = 0.01; interaction: *F*_1, 36_ = 6.01, *P* = 0. 01). In C-E and G-I: statistical analyses using two-way ANOVA followed by Holm-Sidak’s multiple comparisons test. *P* values are provided in the panels. Box plot denotes minimum, first quartile, median, third quartile, and maximum values. Gray lines connecting box plots represent paired data. * *P* < 0.05; ** *P* < 0.01; *** *P* < 0.001; **** *P* < 0.0001.

First, we confirmed that social isolation for 3 or 6 hours (similar to **Fig 1**), but not for only 10 minutes, increases social investigation in juvenile mice (**Figure S1**), similar to previous reports^2,27-29^. To directly test whether Agrp neurons are necessary for the effect of social isolation on increased sociability, we measured social investigation in mice after isolation while chemogenetically inhibiting (with the inhibitory receptor hM4Di) Agrp neurons in P25 (juvenile) and P75 (adult) mice (**Fig. 2B**). We injected the arcuate nucleus of the hypothalamus of newborn Agrp-*IRES*-Cre mice with a viral vector that expresses hM4Di exclusively in Agrp neurons (AAV-DIO-hM4Di-mCherry)^32,33^ (**Figure S2**).

At P25 or P75, we quantified social investigation before and after social isolation and with or without inhibition of Agrp neurons during the period of social isolation. Mice were first tested for their baseline social investigation. After the baseline stage, *Agrp*^hM4Di^ mice were randomly assigned to receive an injection of vehicle (saline) or C21 and then were socially isolated in a new cage. After three hours of social isolation, mice were tested again. Both P25 and P75 control mice display increased social investigation (**Fig. 2C-E** and **Figure S2**). In contrast, chemogenetic inhibition of Agrp neurons blunts the effect of social isolation on social investigation in P25 mice, but not in adults (**Fig. 2C-E**). These findings reveal that Agrp neuron activity is required for the enhanced sociability observed in juvenile but not adult mice following social isolation, indicating that the neural mechanisms regulating social needs are age-dependent.

### Activation of Agrp neurons is sufficient to enhance sociability in juveniles

Considering the aforementioned results, we reasoned that the activation of Agrp neurons may simulate an extended period of social isolation, subsequently promoting social behavior in young mice (**Fig. 2F**). To test this possibility, we quantified social investigation in P25 and P75 mice upon chemogenetic activation of Agrp neurons. We used *Agrp*^Trpv1^ mice^34^ bred in a Trpv1 knockout background, which allow rapid, specific activation (within minutes) of Agrp neurons upon injection of the Trpv1 ligand, capsaicin (see Methods). After measuring baseline social investigation, all mice were injected with capsaicin and isolated in a new cage for five minutes to allow for a robust activation of Agrp neurons in *Agrp*^Trpv1^ mice^34^, before tested again for social investigation (**Fig. 2F**). The social investigation of control mice did not change from the baseline to the test period in both P25 and P75 mice (**Fig. 2G-I**), confirming that five minutes of social isolation does not increase social investigation (**Figure S1**). However, in contrast to control mice, activation of Agrp neurons in P25 mice but not P75 induced a strong increase in social investigation (**Fig. 2G-I**). We further corroborated these findings by activating Agrp neurons using optogenetics, without a previous period of social isolation. In this experiment, again, the activation of Agrp neurons rapidly increased sociability in juveniles (**Figure S3**). Together, these behavioral experiments underscore the critical role of Agrp neurons in promoting sociability in young but not adult mice.

### Social interactions reduce Agrp neuron activity during adolescence, but not in adulthood

If Agrp neurons signal the need for socialization at young ages, then it is predicted that social interactions should reduce the activity of Agrp neurons in socially-deprived mice^20^. To test this idea, we next recorded the activity of Agrp neurons during social isolation and reunion. To isolate the effects of social stimuli and eliminate potential confounding factors, such as multisensory cues from the home cage, we conducted recordings in a neutral cage environment, free from soiled nesting materials or familiar olfactory cues. To record the activity of Agrp neurons, we injected the arcuate nucleus of the hypothalamus of newborn Agrp-IRES-Cre mice^35^ with a viral vector that expresses the calcium sensor jGCaMP7s^36^ (AAV-DIO-jGCaMP7s) exclusively in Agrp neurons. Using a fiber optic probe inserted just above the arcuate nucleus, we assessed neuronal activity by recording jGCaMP7s fluorescence (**Fig. 3A**).

**Figure 3:**
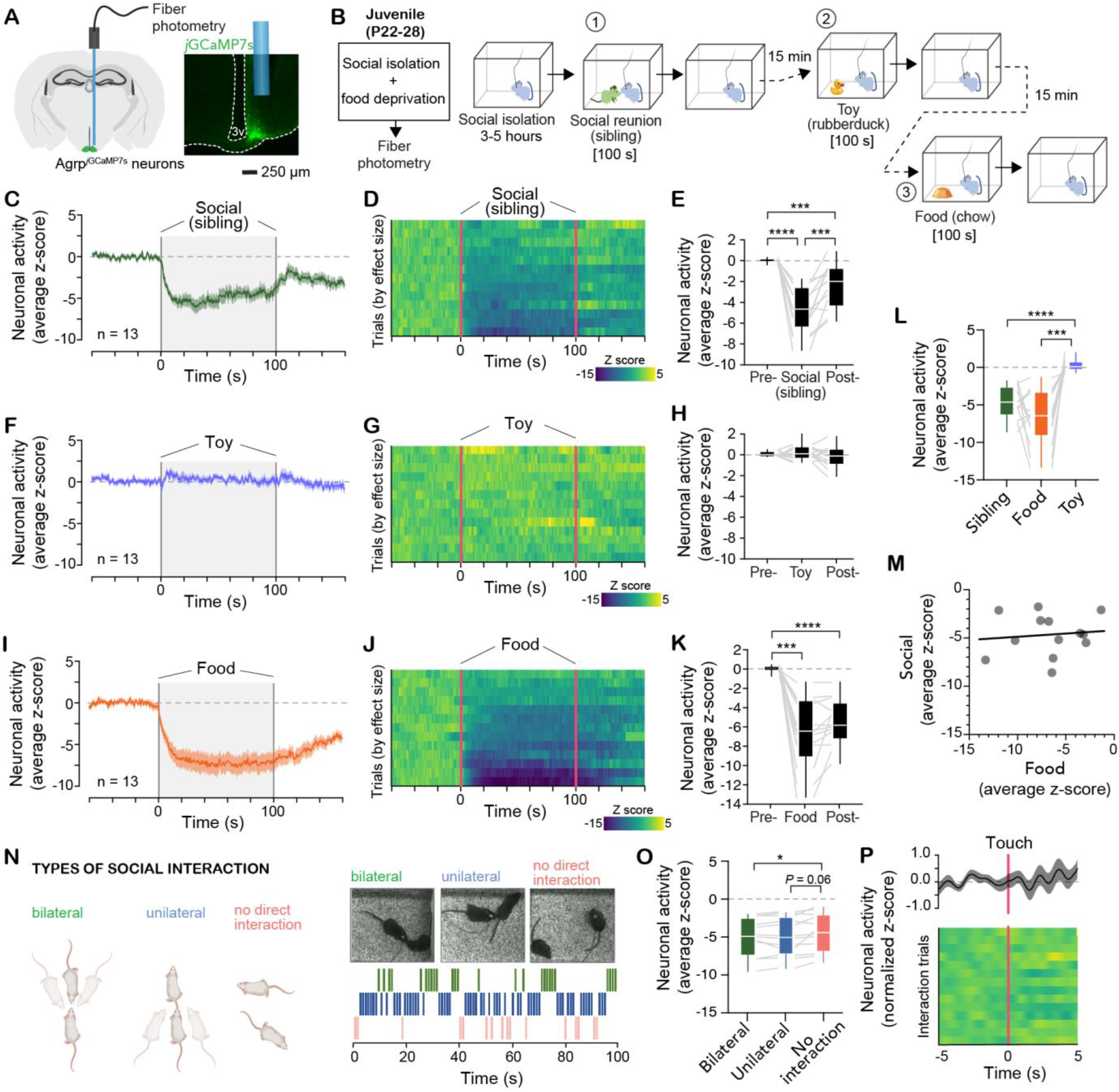
Social interactions reduce Agrp neuron activity in juvenile mice. **(A)** Representative images of fiber optic cannula placement in Agrp^jGCaMP7s^ mice. **(B)** Juvenile mice (P22-P28) were socially isolated, and food deprived for 3-5 hours, then allowed to interact with various stimuli (siblings, toy, food) for 100 seconds. Stimuli were presented sequentially, separated by 15-minute breaks. **(C)** The average Z score of Agrp neuron activity; shaded area represents the period during which the sibling was present in the recording chamber. **(D)** Heatmap showing individual responses (z-score) of Agrp neurons during reunion with a sibling ordered by effect size. **(E)** The mean Z score during baseline, social interaction, and recovery (ε = 0.83, *F*_1.66, 20_ = 37.24, *P* < 10^-6^). **(F-H)** Similar to C-E, but in response to interactions with an inanimate toy control (ε = 0.83, *F*_1.87, 22.52_ = 1.76, *P* = 0.19). **(I-K)** Similar to C-E, but in response to food presentation (ε = 0.64, *F*_1.28, 15.38_ = 42.96, *P* < 10^-5^). **(L)** Comparison of the mean response to sibling, food, and toy (ε = 0.72, *F*_1.44, 17.39_ = 26.57, *P* < 10^-4^). **(M)** Linear regression correlating the mean effect to food with the mean effect to social (*r*^2^ = 0.006, *P* = 0.79). **(N)** Classification of social interaction types based on pose estimation and an example ethogram of behavior classification. **(O)** The mean Z score during the three types of social interaction (ε = 0.75, *F*_1.51, 16.67_ = 6.58, *P* = 0.01). **(P)** Effect of touch on the activity of Agrp neurons (top, average z-score; bottom, heatmap of individual contact trials). In C-M, data corresponds to 13 trials from 9 mice; 4 mice were recorded at two different ages. In E, H, K, L, O: repeated measures one-way ANOVA with Geisser-Greenhouse correction followed by Holm-Sidak’s multiple comparisons test. In C, F, I, P: lines represent mean ± sem. In M, symbols represent individual values. Box plot denotes minimum, first quartile, median, third quartile, and maximum values. Gray lines connecting box plots represent paired data. * *P* < 0.05; ** *P* < 0.01; *** *P* < 0.001; **** *P* < 0.0001.

Mice were socially isolated and food deprived for at least 3 hours (similar to **Figs 1-2**) to monitor the responses to both social reunion and food presentation (**Fig. 3B**). In juveniles (P22-P28) separated from the home cage, reunion with a social stimulus—their same cage, same age, sibling—caused a rapid reduction of Agrp neuron activity that persisted during the entire reunion period (**Fig. 3C-D**). This effect was long-lasting— removal of the sibling from the chamber after the reunion trial did not immediately return the activity of Agrp neurons to baseline levels (**Fig. 3C-E**)—and occurred equally in males and females (**Figure S4**), but not in response to interaction with a control toy (**Figs. 3F-H**). Moreover, as in adults^16,18,19^, food reduced Agrp neuron activity (**Fig. 3I-K**), an effect of similar magnitude as (**Fig. 3L**), but not correlated to (**Fig. 3M**), the response to social reunion, suggesting that these responses are not interdependent.

To determine whether specific aspects of the social interaction between juvenile mice modulate the activity of Agrp neurons, we tracked their social behaviors while recording neuronal activity. Both test and stimulus mice were tracked concurrently, and social interactions were classified based on the distance and angle between the snout of the test mouse and the stimulus mouse^37^ (**Fig. 3N**). The reduction of Agrp neuron activity in the presence of a sibling was greater during active social interactions compared to periods when the mice were not interacting—the mean difference between active interactions and no interactions was -0.49 z-scores (95% CI [-0.85, -0.13]). However, the type of social interaction (unilateral versus bilateral) did not affect the magnitude of this response (**Fig. 3O**). Additionally, Agrp neuron activity did not change when mice physically touched (**Fig. 3P**). These results demonstrate that while the presence of a stimulus mouse in the environment is sufficient to reduce Agrp neuron activity, active social interactions further reduce activity, but this does not necessitate direct contact between mice.

### Social reunion with select conspecifics reduces Agrp neuronal activity in juveniles

Next, we explored whether the reduction in Agrp neuron activity observed during reunion with siblings extends to other social contexts. First, we tested the effect of reunion with peers—unfamiliar mice of the same age. We performed these experiments using all possible sex combinations: male test mice reunited with male or female peers and female test mice reunited with male or female peers. Similar to reunion with siblings, and regardless of sex of the interaction pair, reunion with peers reduced the activity of Agrp neurons (**Fig. 4A-G**; two-way analysis of variance with sex of the test mouse and sex of the stimulus mouse did not reveal any effects of sex). These results support previous reports showing that social isolation increases rebound social investigation in juvenile mice independently of the sex of the interaction pair^2^, and support the idea that Agrp neurons broadly signal social need in juveniles.

**Figure 4:**
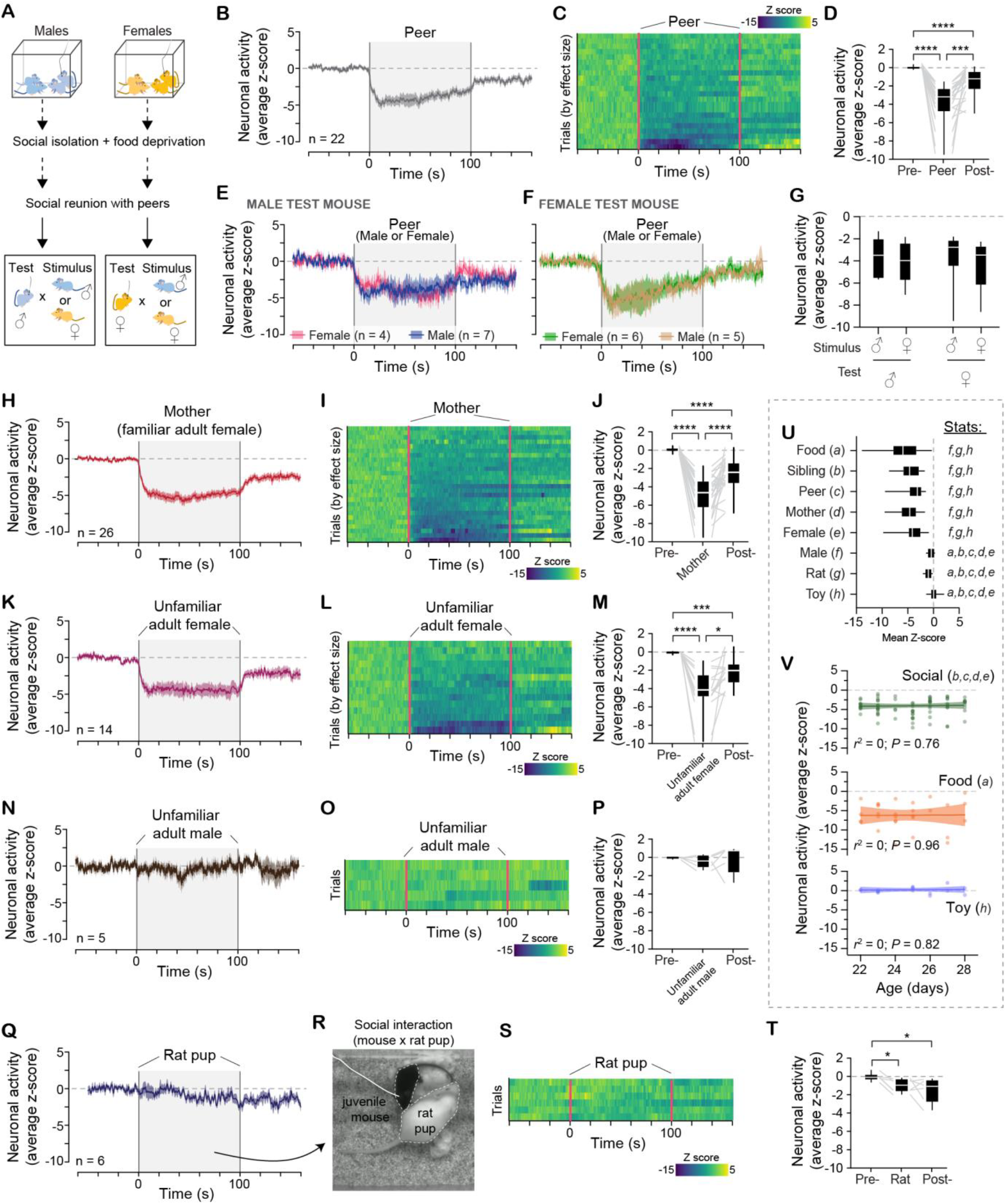
Agrp neurons respond to multiple, but specific, social stimuli. **(A)** Juvenile mice (P22-P28) were socially isolated, and food deprived for 3-5 hours, then allowed to interact with same- or opposite-sex peers for 100 seconds. **(B)** The average Z score of Agrp neuron activity; shaded area represents the period during which a peer—novel mouse of the same age as the test mouse—was present in the recording chamber (data from all sex combinations combined). **(C)** Heatmap showing individual responses (z-score) of Agrp neurons during reunion with a peer ordered by effect size. **(D)** The mean Z score during baseline, social interaction, and recovery (ε = 0.77, *F*_1.54, 32.44_ = 37.98, *P* < 10^-7^). **(E)** The average Z score of Agrp neuron activity of juvenile male mice reunited with same- or opposite-sex peers. **(F)** Similar to E, but for juvenile female mice. **(G)** Comparison of the mean response of male and female mice to same- or opposite-sex peers (effect of sex of test mice: *F*_1, 18_ = 0.13, *P* = 0.71; effect of sex of stimulus mice: *F*_1, 18_ = 0.01, *P* = 0.91; interaction: *F*_1, 18_ = 0.14, *P* = 0.7). **(H-J)** Similar to B-D, but in response to reunion with the mother (ε = 0.90, *F*_1.80, 45.10_ = 73.70, *P* < 10^-13^). **(K-M)** Similar to B-D, but in response to reunion with an unfamiliar adult female (ε = 0.75, *F*_1.51, 19.72_ = 22.91, *P* < 10^-4^). **(N-P)** Similar to B-D, but in response to reunion with an unfamiliar adult male (ε = 0.63, *F*_1.26, 5.06_ = 0.22, *P* = 0.7). **(Q-T)** Similar to B-D, but in response to reunion with a rat pup (ε = 0.66, *F*_1.33, 6.68_ = 6.62, *P* = 0.03). In R, frame depicting active social interaction between the juvenile test mouse and the rat pup. **(U)** Comparison of the mean response to all stimuli tested (*F*_7, 94.81_ = 19.39, *P* < 10^-14^). **(V)** Linear regression correlating the mean Z score during stimulus presentation and age of the mice. In D, J, M, P, T: repeated measures one-way ANOVA with Geisser-Greenhouse correction followed by Holm-Sidak’s multiple comparisons test. In G: two-way ANOVA with sex of test mice and sex of stimulus mice as factors. In U: Brown-Forsythe ANOVA test followed by Dunnett’s T3 multiple comparisons test; letters *a-h* represent pairwise differences between groups (*P* < 0.01 for all comparisons). In B, E, F, H, K, N, Q: lines represent mean ± sem. Box plot denotes minimum, first quartile, median, third quartile, and maximum values. Gray lines connecting box plots represent paired data. In V, symbols represent individual values and lines represent linear regression ± 95% CI bands. * *P* < 0.05; ** *P* < 0.01; *** *P* < 0.001; **** *P* < 0.0001.

To further characterize the response of Agrp neurons to conspecifics, we tested the effect of other familiar and unfamiliar mice: the mother (familiar adult female, which they were separated from P21), unfamiliar adult females, or unfamiliar adult males. We found that social interactions with their mother (**Fig. 4H-J**) and unfamiliar adult females (**Fig. 4K-M**) strongly reduced Agrp neuron activity, while interactions with adult males did not (**Fig. 4N-P**). Moreover, the reduction in Agrp neuron activity is species-specific, as social reunion with an amicable rat pup (of body size similar to an adult mouse) did not elicit the same robust decrease in Agrp neuron activity (**Fig. 4Q-T**).

When comparing the response of Agrp neurons across all stimuli tested, we found that social interactions with siblings, peers, the mother, and unfamiliar adult females reduced Agrp neuron activity to a similar extent; this effect was comparable in magnitude to the response elicited by food presentation (**Fig. 4U**). In contrast, interactions with adult males (although conspecifics), rat pups (non-conspecifics), or exposure to a novel toy did not elicit the same strong reduction in Agrp neuron activity (**Fig. 4U**). Furthermore, the strong response of Agrp neurons to social stimuli was observed at all ages tested, from P22 to P28 (**Fig. 4V**), and occurred even when mice were socially isolated with ad libitum food in the home cage (**Figure S5**). These findings suggest that while social experiences broadly regulate Agrp neuron activity, this modulation is specific to socially significant interactions with members of the same species.

### Olfaction is required for the effects of social reunion on Agrp neurons

Our previous experiments demonstrate that the activity of Agrp neurons reduces in the presence of conspecifics even when the animals do not touch, suggesting that that visual, auditory, or olfactory cues may account for this effect. We designed the next set of experiments to identify which sensory modalities are important for the reduction in Agrp neuron activity upon social interactions in juvenile mice.

We first tested the role of visual processing on Agrp neuron activity by examining the effects of social reunion in darkness (**Fig. 5A**). Within the first ten seconds after reunion, the drop in Agrp neuron activity was similar when mice were tested in light or dark conditions (**Fig. 5B-C**), suggesting that visual inputs are not required for this initial reduction in neuron activity. However, during the remaining period of reunion the reduction in Agrp neuron activity was more pronounced in light compared to dark (**Fig. 5B-D**), suggesting that visual cues may contribute to maintaining the reduced activity of Agrp neurons during social interactions. Thus, while visual cues provide sensory information for the sustained effects of social stimuli on the activity of Agrp neurons, they are not required for the initial detection of the social stimuli upon reunion.

**Figure 5:**
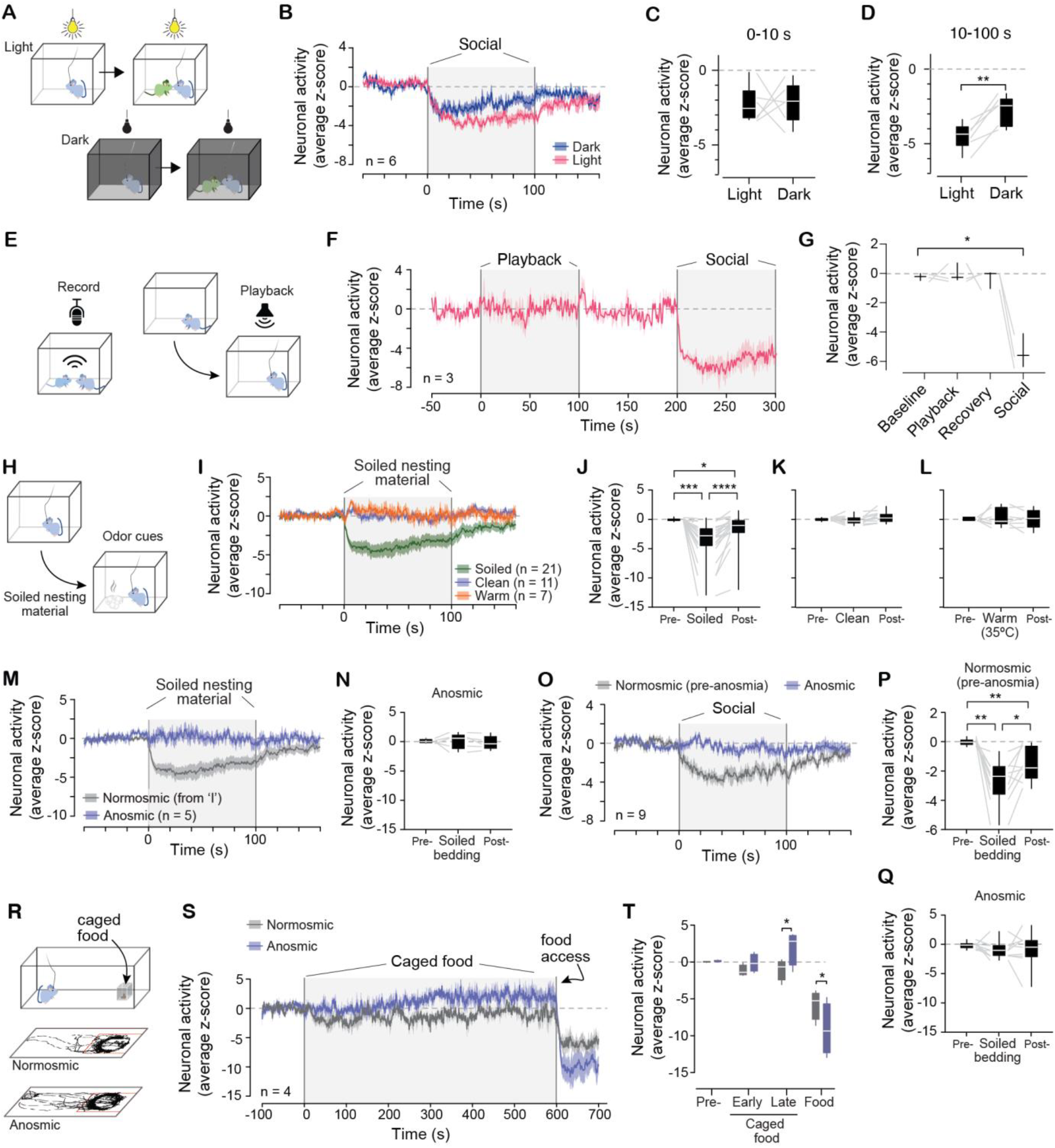
Olfaction is the primary sensory modality involved in Agrp neuron responses to social stimuli. **(A)** Juvenile mice (P22-P28) were tested in light and dark conditions; mice were socially isolated, and food deprived for 3-5 hours, then reunited with a social stimulus mouse (mother). **(B)** The average Z score of Agrp neuron activity; shaded area represents the period during which the social stimulus mouse was introduced to the recording chamber. **(C)** Mean Z scores during the first 10 seconds after reunion (*t*_5_ = 0.10, *P* = 0.92). **(D)** Mean Z scores from 10 to 100 seconds after reunion (*t*_5_ = 5.59, *P* = 0.002). **(E)** Ultrasonic vocalizations were recorded during social interactions between two juvenile mice; these audio recordings were played back to a juvenile mouse while recording the activity of Agrp neurons. **(F)** The average Z score of Agrp neuron activity; shaded areas represent the period during which ultrasonic vocalizations were played back and then the time when the social stimulus mouse was present in the recording chamber as a positive control. **(G)** The mean Z score during baseline, social interaction, and recovery (ε = 0.44, *F*_1.33, 2.67_ = 35.00, *P* = 0.01). **(H)** Protocol to test the effect of odor cues on Agrp neuron activity: juvenile mice were socially isolated and food deprived for 3-5 hours, then presented with nesting material (clean or soiled, at room temperature or warmed to nest temperature—35ºC). **(I)** The average Z score of Agrp neuron activity; shaded area represents the period during which nesting material was presented in the recording chamber. **(J)** Similar to G, the mean Z score during baseline, presentation of soiled nesting material, and recovery (ε = 0.67, *F*_1.30, 26.80_ = 19.99, *P* < 10^-4^). **(K)** Similar to J, but for clean nesting material (ε = 0.75, *F*_1.51, 15.15_ = 3.47, *P* = 0.06). **(L)** Similar to J, but for clean nesting material warmed to 35ºC (ε = 0.69, *F*_1.38, 8.30_ = 0.21, *P* = 0.73). **(M)** Similar experiment as in H-I but testing the effect of soiled nesting material in anosmic mice (ablation of the olfactory epithelium by treatment with methimazole). Normosmic data from panel I. **(N)** Similar to J, but in anosmic mice (ε = 0.64, *F*_1.28, 5.14_ = 0.26, *P* = 0.68). **(O)** The average Z score of Agrp neuron activity of the same mice before (normosmic) and after ablation of the olfactory epithelium (anosmic) in response to social reunion; shaded area represents the time of social reunion. **(P)** Similar to G, mean Z score during baseline, social interaction, and recovery in normosmic mice (ε = 0.82, *F*_1.65, 13.20_ = 18.76, *P* = 0.0002). **(Q)** Similar to P, but for anosmic mice (ε = 0.65, *F*_1.31, 10.48_ = 0.65, *P* = 0.47). **(R)** Normosmic and anosmic mice were tested for their response to a caged food pellet; examples of behavior tracking showing exploration of the caged food. **(S)** The average Z score of Agrp neuron activity of normosmic and anosmic mice; shaded area represents the period during which caged food (inaccessible) was present in the recording chamber. **(T)** Mean Z scores from S during the baseline period, soon after introduction of caged food (i.e., the first 100 seconds, ‘early’), at the end of the exposure to caged food (last 100 seconds, ‘late’), and after uncaging the food (‘food’); effect of time: *F*_3, 9_ = 36.40, *P* < 10^-4^; effect of olfactory status: *F*_1, 3_ = 0.05, *P* = 0.83; interaction: *F*_3, 9_ = 8.43, *P* = 0.005. In C, D: paired *t* test. In G, J, K, L, N, P, Q: repeated measures ANOVA was followed by Holm-Sidak’s multiple comparisons test. In T: two-way ANOVA with time and olfactory status as repeated measure factors. In B, F, I, M, O, S: lines represent mean ± sem. Box plot denotes minimum, first quartile, median, third quartile, and maximum values. Gray lines connecting box plots represent paired data. * *P* < 0.05; ** *P* < 0.01; *** *P* < 0.001; **** *P* < 0.0001.

We next examined the role of auditory cues in modulating Agrp neuron activity, as juvenile mice emit ultrasonic vocalizations during social interactions and these vocalizations are thought to signal a positive affective state, facilitating social interactions^27^. We first recorded the emission of ultrasonic vocalizations during social interactions in juvenile mice (**Fig. 5E**). Next, to test the effect of these ultrasonic vocalizations on the activity of Agrp neurons and simulate auditory cues of conspecifics, we played back these ultrasonic vocalizations to juvenile mice while recording Agrp neuron activity (**Fig. 5E**). Despite these vocalizations’ role in social behavior^27^, their playback did not significantly affect Agrp neuron activity, though the subsequent introduction of social stimulus as a positive control did (**Fig. 5F-G**). These results suggest that auditory signaling does not play a major role in modulating the activity of Agrp neurons in response to social interactions.

We next tested whether olfactory cues are sufficient to reduce the activity of Agrp neurons (**Fig. 5H**). In socially isolated mice, the introduction of soiled nesting material strongly reduced the activity of Agrp neurons (**Fig. 5I-J**), an effect that was not observed with clean nesting material, even when warmed to ≈35ºC (**Fig. 5I, K-L**). To further investigate the effects of olfaction on the activity of Agrp neurons, we rendered mice anosmic by injecting methimazole, a drug that produces extensive degeneration of the olfactory epithelium and subsequent anosmia (see Methods). Introduction of soiled nesting material to methimazole-treated mice had no effect on the activity of Agrp neurons (**Fig. 5M-N**). Furthermore, in anosmic mice, social reunion failed to reduce the activity of Agrp neurons even though all other senses were intact (**Fig. 5O-Q**). These experiments demonstrate that olfaction is the primary sensory modality required for the effects of social stimuli on Agrp neurons.

Notably, in the presence of a caged (and unreachable) food pellet, when anosmic mice approach and explore the caged food (**Fig. 5R**), the activity of Agrp neurons increased, instead of decreasing (**Fig. 5S-T**). As soon as the food pellet is uncaged, when mice bite and ingest food, we observed a robust reduction in the activity of Agrp neurons (**Fig. 5S-T**). These results demonstrate that olfaction is critical for Agrp neuron responses to both social and food stimuli. However, in contrast to eating, olfaction completely abrogates responsivity to social stimuli whereas Agrp neurons are still responsive to the consummatory aspects of eating. Together, these experiments demonstrate a broad role for olfactory signaling in the modulation of Agrp neurons: in the absence of olfaction, only food consumption, but not food detection or social interaction, reduces the activity of Agrp neurons.

### The response of Agrp neurons to social stimuli diminishes with age

When do Agrp neurons stop responding to social stimuli? To address this question, we recorded Agrp neuron activity from P30 through adulthood (see **Fig. 1A**). Mice were both socially isolated and food deprived to ensure robust activation of Agrp neurons across all ages and to compare their response to different social stimuli, food, and toy. Similar to our results in juvenile mice, we found that the reduction in Agrp neuron activity upon reunion with siblings (**Fig. 6A-B**), peers (**Fig. 6C-D**), mother (**Fig. 6E-F**), and unfamiliar adult females (**Fig. 6G-H**), but not a toy control (**Fig. 6I-J**), gradually decreased in intensity as mice age. The lack of response to social interactions in older mice was not due to an overall decrease in Agrp neuron responsiveness to sensory stimuli, as the presentation of food reduced Agrp neuron activity to a similar extent across this entire age range (**Fig. 6K-L**). Social reunion failed to alter Agrp neuron activity after 65 days of age, even when mice were socially isolated and food deprived overnight (**Figure S6**). Altogether, these results demonstrate that Agrp neuron activity in response to social isolation and reunion is restricted to the developmental window from infancy to late adolescence.

**Figure 6:**
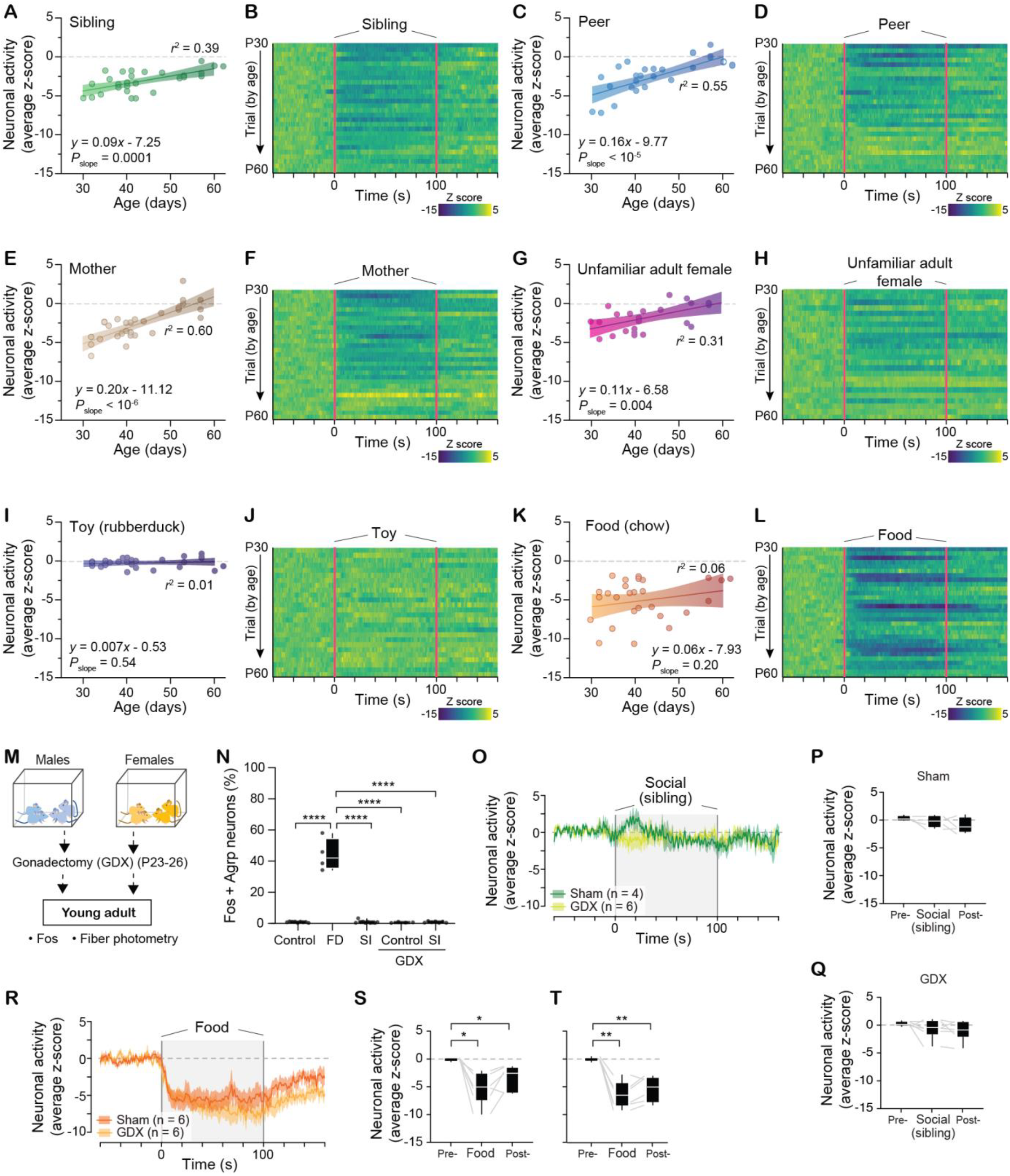
The reduction of Agrp neuron activity by social interaction slowly disappears during adolescence. **(A-L)** The activity of Agrp neurons was recorded using fiber photometry in mice from P30-60, covering the period of adolescence (see Fig. 1A). After mice have been separated from the home cage for 3-5 hours, different stimuli were introduced to the recording chamber: sibling (A-B), peer (mouse at the same age from a different cage, C-D), mother (E-F), unfamiliar adult female (a non-lactating adult female unknown to the experimental mouse, G-H), a control toy (I-K), and food (pellet of chow, K-L). **(A)** Linear regression correlating the mean Z score during reunion with sibling and age of the mice (n = 28 trials from 10 mice). **(B)** Heatmap showing individual responses (z-score) of Agrp neurons during reunion with a sibling ordered by age of recordings. **(C-D)** Similar to A-B, but for peers (n = 28 trials from 9 mice). **(E-F)** Similar to A-B, but for the mother (n = 29 trials from 10 mice). **(G-H)** Similar to A-B, but for unfamiliar adult female (n = 24 trials from 8 mice). **(I-J)** Similar to A-B, but for control toy (n = 26 trials from 9 mice). **(K-L)** Similar to A-B, but for food (n = 30 trials from 12 mice). **(M)** Juvenile mice (P23-P26) were gonadectomized; when young adults, mice were socially isolated (SI) and/or food deprived (FD) overnight to quantify activation of Agrp neurons with Fos and the response of Agrp neurons to social/food stimuli. **(N)** Agrp neurons positive for Fos (control, n = 9; FD, n = 4; SI, n = 10; GDX-control, n = 4; GDX-SI, n = 7; *F*_4, 29_ = 143.46, *P* < 10^-14^). **(O)** The average Z score of Agrp neuron activity in control (sham) and GDX mice; shaded area represents the period during which a sibling was introduced to the recording chamber. **(P)** The mean Z score during baseline, social interaction, and recovery for sham mice (ε = 0.77, *F*_1.55, 4.67_ = 2.04, *P* = 0.22). **(Q)** Similar to P, but for GDX mice (ε = 0.79, *F*_1.59, 7.99_ = 2.03, *P* = 0.19). **(R-T)** Similar to O-Q, but in response to presentation of food: sham control (ε = 0.80, *F*_1.60, 8.04_ = 13.84, *P* = 0.003); GDX (ε = 0.63, *F*_1.27, 6.37_ = 36.19, *P* = 0.0006). In N: one-way ANOVA. In P, Q, S, T: repeated measures one-way ANOVA with Geisser-Greenhouse correction. ANOVA was followed by Holm-Sidak’s multiple comparisons test. In A, C, E, G, I, K: symbols represent mean Z-score from individual recordings; lines represent linear regression ± 95% confidence interval; *r* square and *P* value for the linear regression analysis are shown in the panels. In O, R: lines represent mean ± sem. In P, Q, S, T, box plot denotes minimum, first quartile, median, third quartile, and maximum values and gray lines connecting box plots represent paired data. * *P* < 0.05; ** *P* < 0.01; *** *P* < 0.001; **** *P* < 0.0001.

Given the importance of gonadal hormones during adolescence for neurocircuitry and behavioral maturation, we next tested whether the rise of these hormones during adolescence^26^ contributes causally to the decline in the responsivity of Agrp neurons to social contexts. To test this, we surgically removed the gonads of male and female mice before puberty, thereby preventing the surge in gonadal hormones during adolescence, and measured Agrp neuron response to social isolation and reunion in young adulthood (**Fig. 6M**). In gonadectomized mice (both male and female), social isolation failed to activate Agrp neurons (**Fig. 6N**). Moreover, social interactions failed to reduce Agrp neuron activity in animals socially isolated and food deprived (**Fig. 6O-P**), while presentation of food reduced Agrp neuron activity in both groups of mice (**Fig. 6R-T**). Thus, the decline in responsiveness of Agrp neurons to social stimuli during the transition to adulthood occurs independently of the emergence of gonadal hormones during adolescence

## Discussion

### Summary

This study reveals that the canonical ‘hunger’ neurons in the brain (Agrp neurons) coordinate divergent need states and behaviors at different periods of life. During a sensitive period of development that spans the entire adolescence in mice, social isolation activates, while reunion reduces, the activity of Agrp neurons. These activity changes are important for the modulation of social behaviors. In young mice, but not in adults, the inhibition of these neurons blunts the sociability effects of social isolation, while their activation increases social investigation. These findings demonstrate that a neuron population known to coordinate the need for food in adults is required for proper socialization at younger ages.

### Changes in neuron responses during development

Overall, the influence of social isolation and reunion on Agrp neuronal activity gradually diminishes and eventually disappears following late adolescence. Coinciding with this change, increased agonistic, territorial, and socio-sexual behaviors towards conspecifics begin to manifest during adolescence, implying that Agrp neurons contribute to the neural mechanisms regulating infantile and adolescent social behaviors prior to the emergence of sex-specific social behaviors in early adulthood. While our results rule out gonadal hormones as primary mediators of the transition in Agrp neuron responses from adolescence to adulthood, the extent to which changes in other neuroendocrine systems affect the development of Agrp neurons and neuronal circuits involved in social behaviors is an important point for future research.

Following a period of social isolation, mice exhibit a preference for contexts associated with social interactions over those linked to social isolation^27,28^. This preference peaks around adolescence and declines in early adulthood^38^, similar to our results. This shift in preference relies on oxytocin signaling within the ventral striatum^38^. During social interactions, oxytocin from the hypothalamus modulates both the ventral striatum and the midbrain dopamine neurons that project to the ventral striatum, amplifying the positive value of social interactions^39,40^—a phenomenon more pronounced in younger individuals^38^. These findings, together with the current work showing that Agrp neurons respond to social contexts and affect social behaviors in adolescence but not adulthood, support the idea that specific neural mechanisms differentially influence social behaviors at discrete developmental stages, possibly reflecting changes in the adaptive function of such behaviors.

### Hunger neurons that regulate social homeostasis

The results of this study should be considered alongside the extensive research on the functional properties of Agrp neurons in energy metabolism. Agrp neurons exhibit increased activity during negative energy states^14,15^ and decreased activity when consuming or anticipating a caloric meal^16-19,41^. In young mice, social stimuli parallel the effect of food on Agrp neuron activity: social deprivation increases Agrp neuron activity and social reunion reduces the activity of these neurons. A notable difference, however, is that while food ingestion (or the consummatory phase of eating behavior) further suppresses Agrp neuron activity, the consummatory phase of social interactions—such as touching and huddling— fails to have a similar effect. Thus, the response of Agrp neurons to social interactions is analogous to the anticipatory (preingestive) effect of food on the activity of these neurons. Moreover, because not all social stimuli reduce the activity of Agrp neurons, our results suggest that the age-specific salience of the social stimulus is encoded at the level of Agrp neuron activity.

A recent study identified neurons in the adult mouse preoptic area that regulate social homeostasis. These neurons respond to social isolation and reunion, modulating behaviors associated with social need^42^. Interestingly, this study demonstrated that touch was the critical sensory modality mice used to detect conspecific presence and fulfill their social needs. These findings contrast with our recordings of Agrp neuron activity in juvenile mice, which indicate that olfaction— not touch—is the primary sensory modality mediating the effect of social reunion on Agrp neurons. These findings could be interpreted to suggest that, in young animals, Agrp neurons function to ensure proximity with conspecifics—prioritizing physical closeness over direct social contact. This interpretation aligns with attachment theory, which posits that infants possess an innate attachment system designed to maintain social homeostasis, with the outcome of the activation of this attachment system being proximity to the attachment figures. These proximity-keeping behaviors are foundational for establishing emotional and physical connections in early development^7^. As animals mature, this attachment system evolves, and in adolescence, proximity-keeping broadens to peers, supporting social integration. Our results therefore imply that Agrp neurons are a key population of neurons addressing the social needs of mammals during early development, thus forming part of the long-sought neural network that maintains social homeostasis, as predicted by attachment theory^7^.

### Repurposing of the hunger circuitry for social needs

Evolution can repurpose (or exapt) traits that originally developed for one function to fulfill a new function^43^. Since Agrp neurons in the hypothalamus are phylogenetically older than mammals, the response of these neurons to the social environment during early development in mice lends support to the idea that the feeding circuitry was repurposed during mammalian evolution to address the necessity for robust social bonds during infancy and adolescence. Intriguingly, studies in worms and flies similarly suggest that some neural mechanisms associated with feeding and metabolism also affect social behaviors^44,45^. These observations may indicate that the cognitive requirements for social and ingestive behaviors intersect across taxa^46^, justifying the adaptive value in repurposing neural mechanisms to support both functions.

Nonetheless, this repurposing of neural mechanisms is not confined solely to feeding and social behaviors. Recent research proposes that a specific group of neurons in the mouse brain was repurposed from an ancestral arousal network to facilitate maternal behavior^47^. This conceptual framework is supported by theoretical work, arguing that the brain’s central organizational principle is the reallocation or repurposing of neural mechanisms for various tasks^48^.

In sum, this study provides a neurobiological underpinning for the regulation of social needs during early development^49^ and how these mechanisms change during the complex series of developmental events that occur from infancy to adulthood.

## Acknowledgements

We thank members of the Dietrich lab for critical feedback on the project and on the manuscript. We thank the Janelia GENIE project for the *j*GCaMP7s plasmid. M.O.D. was supported by the National Institute of Mental Health of the National Institutes of Health (R01MH125008 and R01MH130825), by the Charles H. Hood Foundation, Inc. (Boston, MA), by the Smith Family Foundation, by a grant of the Foundation for Prader-Willi Research, by the Reginald and Michiko Spector Award in Neuroscience, and by discretionary funds from the Yale School of Medicine. L.K. was supported by Yale College First-year Summer Research Fellowship and by Yale College Dean’s Research Fellowship. We thank David Bruin for copyediting the manuscript.

## Author contributions

M.O.D and O.I conceived the hypothesis, designed the study, analyzed the data, and wrote the manuscript. O.I, L.K, M.R.Z., J.B., Y.L., and M.P. performed experiments. G.M. developed software to analyze and plot the data. All authors read and edited the manuscript.

## Materials and Methods

### Mice

Both male and female mice were used in the studies. Agrp-IRES-Cre founder mice were obtained from the Jackson Laboratories (Jax #012899). For Fos experiments, Agrp^HA^ mice were generated by breeding Agrp-IRES-Cre mice with Rpl22^LSL-HA^ mice (Jax #029977). Agrp^tdTomato^ mice were generated by breeding Agrp-IRES-Cre mice with Ai14(RCL-tdT) (Jax #007914) or Ai75(RCL-nls tdT) (Jax #025106) for nuclear-localized tdTomato fluorescence. For the rapid activation of Agrp neurons, Agrp^Trpv1^ mice were generated from Agrp-IRES-Cre mice crossed to R26^LSL-Trpv1^ mice that were backcrossed to Trpv1 knockout mice as we have previously characterized^34^. Thus, Agrp^Trpv1^ mice were AgrpCre^Tm/+^::Trpv1^—/—^::R26-LSL-Trpv1^Gt/+^ and littermate control animals were Trpv1^—/—^:R26-LSL-Trpv1^Gt/+^ mice injected with capsaicin^34,50,51^. Analysis of ectopic expression of Cre was performed by using a specific set of primers against the excised conditional allele, as previously reported^34^. Mice exhibiting ectopic expression of the excised allele were excluded from the subsequent studies and not included in the analysis. All mice were maintained in our own colony kept in temperature- and humidity-controlled rooms, in a 12/12 h light/dark cycle, with lights on from 7:00 AM–7:00 PM. Food (Teklad 2018S, Envigo) and water were provided ad libitum. All procedures were approved by Institutional Animal Care & Use Committee (IACUC) of Yale University.

### Drugs

Buprenorphine (0.1 mg/kg dissolved in sterile saline, s.c.). Meloxicam (5 mg/kg in sterile saline, s.c.). DREADD agonist 21 (Compound 21; 3 mg/kg dissolved in sterile saline). Capsaicin (10 mg/kg, s.c. or i.p.; 3.33% Tween-80 in sterile PBS). Methimazole (1-methyl-2-mercaptoimidazole; 50 mg/kg dissolved in sterile water). Mice were injected at 10 μl/g of body weight.

We administered all compounds at a dose of 10 μl/g of body weight. Analgesia before and after surgeries was provided by buprenorphine (0.1 mg/kg dissolved in sterile saline given subcutaneously) and meloxicam (5 mg/kg in sterile saline given in intraperitoneally). DREADD agonist 21 (Compound 21; 3 mg/kg dissolved in sterile saline) was delivered intraperitoneally. Capsaicin (10 mg/kg; 3.33% Tween-80 in sterile PBS) was delivered intraperitoneally to activate Trpv1 expressing Agrp neurons. To induce anosmia, methimazole (1-methyl-2-mercaptoimidazole; 50 mg/kg dissolved in sterile water) was administered intraperitoneally^52-54^.

### Fos studies

#### Experiment reported in Figures 1B-E

Agrp^HA^ mice at postnatal day 21 (P21) or 28 (P28) were randomly assigned to one of two conditions: (1) control group, in which mice remained in the home cage with their littermates and had ad libitum access to food; or (2) social isolation group, in which mice were individually housed in a new cage with fresh bedding and ad libitum food for 90 minutes.

#### Experiment reported in Figures 1F-H

Agrp^Ai75^ mice at P25, P35, P50, or P60+ (adults) were randomly assigned to one of three conditions: (1) control group, in which mice remained in the home cage with their littermates and had ad libitum access to food; (2) social isolation group, in which mice were individually housed in a new cage with fresh bedding and ad libitum food for 4.5 hours; or (3) food-deprived group, in which mice were housed in a new cage in social groups without food for 4.5 hours.

#### Experiment reported in Figure 6N

Agrp^Ai14^ mice between P22 and P24 were gonadectomized (see description of procedures under ‘Gonadectomy’). Control mice were sham-operated; some of the control mice and food-deprived mice were not operated, but because results did not differ between sham-operated and non-operated mice, results were pooled together. Once mice reached adulthood (P75), mice were randomly assigned to one of these groups: (1) control group, in which mice remained in the home cage with their littermates and had ad libitum access to food; (2) food-deprived group, in which mice were housed in social groups without food; and (3) social isolation group, in which mice were individually housed in a new cage with fresh bedding and ad libitum food. Gonadectomized mice were randomly assigned to groups (1) or (3). For this experiment with adult animals, mice were socially isolated or food deprived overnight from 7:00 PM to 11:00 AM.

### Immunohistochemistry and quantification of Fos-positive neurons

Mice were deeply anesthetized using inhaled isoflurane in an induction chamber and subjected to cardiac perfusion using freshly prepared fixative (4% paraformaldehyde in 1x PBS with a pH of 7.4). The brains were carefully removed from skull to avoid damaging the median eminence and the arcuate nucleus of the hypothalamus and post-fixed overnight in the fixative at 4ºC. Brains were then sectioned using a vibratome (Leica, VT100s) and coronal brain sections with a thickness of 50 μm were collected in sequence. Brain sections were first washed multiple times in 1x PBS (pH = 7.4) and pre-treated with a solution of 0.3% Triton X-100 in 1x PBS for a duration of 30 minutes. Next, the sections were incubated in a blocking solution (0.3% Triton X-100, 10% Donkey Serum, and 0.3M Glycine in 1x PBS) for one hour. Following the blocking step the sections were incubated with primary antibodies overnight with: (1) rabbit monoclonal anti-Fos antibodies (1:1000, #2250; Cell Signaling Technology) and mouse polyclonal anti-HA antibodies (1:1000; 901503, Biolegend) for experiments with Agrp^HA^ mice; (2) rat monoclonal anti-Fos antibodies (1:1000, #226017, SYSY) and rabbit polyclonal anti-mCherry antibodies (1:1000, #600401, Rockland) for experiments with Agrp^Ai14^ and Agrp^Ai75^ mice. Subsequently, sections were thoroughly washed with 0.3% Triton X-100 in PBS and then incubated with secondary antibodies (For Agrp^HA^ mice: donkey anti-mouse 488, AB_141607 and donkey anti-rabbit 647, AB_2536183; for Agrp^Ai14^ and Agrp^Ai75^ mice: donkey anti-rabbit 594, A-21207 and goat anti-rat 647, A-21247; fluorescent Alexa antibodies were from Thermofisher and used 1:500). Finally, sections were thoroughly washed with 0.3% Triton X-100 in PBS and mounted on glass slides for imaging. Confocal imaging was used to image samples using a Leica Stellaris 5 Confocal microscope (Department of Comparative Medicine, Yale School of Medicine).

Throughout the entire procedure, the investigators were blinded to the experimental groups. The number of Agrp^HA^ and Agrp^Ai14^ neurons and Fos-positive neurons were manually counted using ImageJ (version 1.51h, NIH, USA); for Agrp^Ai75^ animals, Agrp neurons and Fos expressing double-labeled nuclei were automatically counted using a custom-built macro.

### Surgery: viral injections

Newborn Agrp-IRES-Cre mouse pups (P0 or P1) were individually removed from the home nest and cryo-anesthetized on ice using an aluminum foil barrier for 10 minutes. Subsequently, the pups were transferred to a chilled neonatal frame (Stoelting Co., 51600) mounted on a stereotaxic apparatus, where they were head-fixed with head bars and their skull positions aligned. Viral injections were performed using a Hamilton syringe (32-gauge, 15° bevel; Hamilton Co., 81430) loaded with 0.3 µL of an adeno-associated virus (AAV). The arcuate nucleus was targeted using the following coordinates relative to lambda: AP = +1.0 mm, ML = ±0.3 mm, DV = −4.1 mm). After the injections, pups were removed from the frame, placed on thermal support until recovery from anesthesia, and then returned to the home cage.

For photometry experiments, unilateral injections were performed of virus expressing the calcium sensor jGCaMP7s (AAV8-CAG-FLEX-jGCaMP7s-WPRE-SV40, Penn Vector Core and AAV1-CAG-FLEX-jGCaMP7s-WPRE-SV40, Addgene #104495). For optogenetic experiments, bilateral injections were performed of virus expressing channelrhodopsin (AAV9-Flex-rev-ChR2(H134R)-mCherry, Addgene #18916). For behavior experiments bilateral injections were performed of virus expressing the Gi-coupled hM4Di receptor (AAV8-hSyn-DIO-hM4D(Gi)-mCherry, Addgene #44362).

### Surgery: cannula implantation

Agrp-IRES-Cre mice that had received AAV injections as newborns underwent cannula implantation. Mice were given a subcutaneous injection of buprenorphine (0.1 mg/kg in sterile saline, 10 μl/g body weight) 15–30 minutes prior to surgery. Anesthesia was induced by inhalation of isoflurane (2-3 %) in oxygen (1.5 L per min low rate), and body temperature was maintained at 33°C using a controlled heating pad (WPI, animal temperature controller); eye ointment (Dechra) was applied to protect the eyes. Mice were then secured in a stereotaxic apparatus and, following a small scalp incision to expose the skull, a burr hole was drilled at coordinates relative to bregma (AP = +1.4 mm, ML = ±0.3 mm). For photometry experiments, a fiber-optic cannula (400 µm core, NA 0.48, Doric Lenses) was lowered to a depth (DV) of 5.8 mm, whereas for optogenetic experiments the cannula was lowered to 5.6 mm. The cannula was fixed in place using dental cement, the incision was closed with sterile sutures, and mice were subsequently administered meloxicam intraperitoneally (5 mg/kg in sterile saline, 10 μl/g body weight). Mice were allowed to recover for at least 2 days before testing.

### Surgery: gonadectomy

Agrp^Ai14^ mice for Fos experiments or Agrp^Gcamp7s^ for fiber photometry were gonadectomized between P23-P26. Buprenorphine injection was given (0.1 mg/kg dissolved in sterile saline, s.c., 10 μl/g body weight) 15-30 min before the onset of surgery. Subsequently, anesthesia was induced by inhalation of isoflurane (2-3 %) in oxygen (1.5 L per min low rate). Prior to incision, animals received a subcutaneous injection of bupivacaine along the incision line. Body temperature was maintained at 33°C, using a controlled heating pad (WPI, animal temperature controller). The eyes were protected from drying by eye ointment (Dechra).

#### Orchiectomy

A 1 cm ventral midline incision was made just cranial to the prepuce and the linea alba was incised to enter the peritoneal cavity. The testicular fat pad was located and exteriorized along with the testes and epididymis using blunt forceps. The spermatic cord was clamped with a hemostat and ligated with a 5-0 absorbable suture. The testicle and cord were then excised distal to the ligature. This procedure was repeated on the contralateral side. The muscle layer was then closed with a continuous 4-0 absorbable suture and the skin layer closed with interrupted 5-0 monofilament nylon sutures.

#### Ovariectomy

A 1 cm dorsolateral longitudinal skin incision was made approximately midway between the last rib and the hip. Subcutaneous tissue was gently separated, and a small 3–4 mm incision was made through the underlying muscle. The ovarian fat pad and ovary was pulled through the incision with blunt forceps and hemostat clamped over the oviduct. A 5-0 absorbable ligature was tied beneath the clamp and the ovary was then excised distal to the suture. The tissue stumps were allowed to retract into the peritoneal cavity. These steps were repeated on the opposite side, after which the muscle layer was closed with 4-0 absorbable sutures and the skin incision was closed with 5-0 monofilament nylon sutures.

After surgery, mice were allowed to recover in a clean cage partially placed over a warmed pad and then returned to the homecage. Buprenorphine (0.1 mg/kg, s.c.) was repeated every 12 hours for 48 hours, and meloxicam (5 mg/kg, s.c.) was administered once daily for 48 hours. Incision sites were inspected daily for signs of inflammation and non-absorbable skin sutures were removed 10-14 days postoperatively.

### General procedures for fiber photometry recordings

Agrp^jGCaMP7s^ mice implanted with a fiber-optic cannula for recording Agrp neuron activity were used to assess responses to various stimuli. Prior to each experiment, individual mice were transferred to a testing cage with clean bedding and without food unless otherwise specified. The social isolation period for each animal began between 10:00 AM and 1:00 PM, and recordings started 3 to 5 hours after isolation onset.

### Fiber photometry recordings in juvenile mice

For experiments reported in Figure 3, juvenile mice (between P22 and P28) were used in the experiments. Each recording session started with a 3-minute baseline period, followed by the introduction of a sibling for 100 seconds. After the stimulus was removed, the recording continued for an additional 3 minutes. Following a 15-minute break, the same procedures were followed for the presentation of a toy control (rubber duck). Finally, at the end of the experiment, the same procedures were used for the presentation of food (a pellet of chow diet). A total of 9 mice were recorded for all 3 stimuli (sibling, toy, and food) with 4 mice recorded in two different ages, for a total of 13 trials. Some of the mice were also recorded in response to other social stimuli. For experiments reported in Figure 4, similar procedures were used as described above. Different social stimuli were presented in random order. The following social stimuli were used in these experiments: peers (mice of same age but unfamiliar to the test mouse) of both sexes, mother (their biological mother that they had been separated at P21), unfamiliar adult female (an adult female that is non-lactating and unfamiliar to the test mouse), and unfamiliar adult male. These animals were also tested for a toy control and for food presentation. Because of the number of stimuli, not all stimuli were presented at the same day. Some mice were recorded at multiple ages.

For experiments using a rat pup, we followed an identical recording protocol—3 min baseline, 100 s interaction, 3 min post-stimulus—except that, in place of a mouse conspecific, each juvenile mouse was paired with rat pup (Wistar strain, P13–P16). Prior to each recording, mice were isolated for 3 h in a fresh cage as described above.

### Fiber photometry recordings in juvenile mice: sensory modalities

To investigate whether visual inputs contribute to the Agrp neuron response to social stimuli, juvenile mice were first isolated for 3 hours (as described above). Thirty minutes before the testing session, the lights were turned off for mice to habituate to dark conditions. Following this adaptation period, animals underwent a fiber photometry recording protocol identical to the one described above: a 3-minute baseline, followed by a 100-second exposure to social stimuli, and a final 3-minute post-stimulus recording period.

For the experiment investigating the effect of auditory cues, we first recorded ultrasonic vocalizations (USVs) during the reunion of juvenile siblings after a period of 3 hours of social isolation. These recordings were obtained using the UltraSoundGate 416H recorder module in coupled with the CM16/CMPA condenser ultrasound microphone (Avisoft Bioacoustics, Berlin, Germany), which was positioned 15 cm above the animal. Subsequently, we synthetized a condensed audio file containing 100 seconds of these recorded USVs. During fiber photometry recordings, juvenile mice first underwent a 3-minute baseline period, followed by playback of the synthesized USVs for 100 seconds, and then an additional 100 seconds of recording post-playback. Immediately afterward, a social stimulus was presented for another 100 seconds to serve as a positive control. Ultrasonic playback was delivered through Electrostatic speaker ESS16 (Avisoft Bioacoustics, Berlin, Germany).

For experiments testing the effect of nesting material, three different stimuli were used. First, soiled nesting material was collected from the cage of experimental animal. Second, clean nesting material was used by manually shredding a pellet of pressed cotton nesting material using clean gloves to mimic the natural state of the nesting material in the home cage. Third, this clean nesting material was incubated in a warm chamber until it reached 35°C. For experiments involving the presentation of warm nesting material, a thermocamera (Xi 400, Optris Infrared Sensing) was used to monitor the temperature of the nesting material during fiber photometry recordings. The same protocol for stimulus presentation as reported above was used in these studies.

For experiments with anosmic mice, baseline photometry responses were recorded in Agrp^jGCaMP7s^ animals at P22 or P23. After these recordings animals were treated with methimazole (50 mg/kg, i.p.) to cause tissue-selective toxicity of the olfactory mucosa^52-54^. Two days after treatment with methimazole, mice were tested again. Anosmic mice were tested in response to soiled nesting material, social stimuli (mother), and food.

### Fiber photometry recordings: progression from adolescence to adulthood

For experiments reported in Figure 6, assessing the transition from adolescence (P30) to adulthood (P60), were conducted fiber photometry recordings using similar procedures as described above. Different social (siblings, peers, mothers, unfamiliar adult females) and control (toy, food) stimuli were presented in random order. The same mice were recorded at multiple ages.

### Fiber photometry recordings: gonadectomized mice

Fiber photometry recordings were performed using a dual-wavelength excitation scheme to monitor jGCaMP7s fluorescence. A 460 nm LED (calcium-dependent channel) was sinusoidally modulated at 531 Hz, and a 405 nm LED (isosbestic control channel) at 211 Hz. The two beams were combined in a fiber-coupled minicube (FMC4, Doric Systems) and delivered to the implant via a 400 µm-core, 0.48 NA patch cord. Emitted light was collected back through the same path, separated in the minicube, and detected on a femtowatt photoreceiver (Model 2151, Newport; gain set to DC-low). The analog outputs were sent to an RZ5P real-time processor (Tucker-Davis Technologies), where each excitation frequency was demodulated by lock-in amplification. Demultiplexed signals were exported to MATLAB for offline analysis. To correct for motion and photobleaching artifacts, the 405 nm control trace was fitted to the 460 nm signal via polynomial regression; ΔF/F was then computed as (460 nm signal – 405 nm fitted signal) / 405 nm fitted signal. The resulting ΔF/F trace was low-pass filtered to remove high-frequency noise and z-scored relative to the mean and standard deviation of a 60-second pre-stimulus baseline. Fiber photometry recordings and behavior data were synchronized with a TTL signal from Tucker-Davis Technologies software.

### Behavior classification

To classify active and passive social interactions between juvenile mice, we manually tracked frame-by-frame the position of the two mice during our recordings. Behaviors were classified into three categories according to the perspective of the pup as reported^37^. For unilateral interactions, the snout of the experimental mice was interacting with the body of the social stimuli, while the snout of the social stimuli did not interact with the experimental animal. For bilateral interactions, both snouts were classified as interacting with the other animal. All frames in which the snout of the experimental animal was not interacting with the social stimuli were classified as non-interacting. An interaction event was classified if the distance was less than a threshold of 5 cm and within an angle between -110° to +110°, representing the visual field of the mice.

### Sociability testing

All behavioral assays were conducted during the light phase (1 PM to 6 PM) under uniform, low-level white illumination. The three-chamber apparatus (65 × 42 × 23 cm) was constructed of clear red plexiglass and divided into three equally sized compartments by opaque partitions. The center chamber served as a neutral zone; the two side chambers each contained a 10 cm-diameter wire cup that held either the social stimulus (an unfamiliar, age-, sex-, and strain-matched mouse) or a novel object (a small plastic toy). Social and non-social stimuli were randomly assigned to left or right chambers and counterbalanced across subjects. Test mice were allowed free access to the entire arena. Arenas and cups were cleaned between trials with 75% ethanol, rinsed with water, and dried with paper towels to eliminate olfactory cues.

Animal position and head orientation were tracked at 30 Hz using Any-Maze (Stoelting Co.; RRID:SCR_002798). “Social investigation” was defined as time spent with the head of the test mouse within 5.5 cm from the cup containing the stimulus mouse; object investigation was scored identically. Each session of social investigation lasted 15 minutes.

#### Experiment testing the effect of different periods of social isolation on social investigation

At postnatal day 25 (P25), each test mouse first underwent a 15-minute baseline sociability test. Immediately afterward, both the test mouse and its designated social stimulus mouse were singly housed for one of three isolation intervals (15 min, 3 h, or 6 h). Following isolation, mice were returned to the three-chamber arena for a second 15-minute sociability test. This within-subject design allowed direct comparison of social investigation before and after varied social isolation periods.

#### Experiment testing the effect of Agrp neuron inhibition on social investigation

Agrp–IRES–Cre mice expressing the inhibitory DREADD hM4Di were first recorded during an initial 15-minute baseline sociability test. Immediately afterward, mice were injected with Compound 21 (3 mg/kg, i.p., in sterile saline) and singly housed for 3 hours. The arena was cleaned as above during this interval. Following isolation, each mouse underwent a second 15-minute sociability test with the same social and object stimuli.

#### Experiment testing the effect of Agrp neuron activation on social investigation

Agrp^Trpv1^ and littermate control mice were tested as above for a 15-minute baseline. Fifteen minutes into the trial, mice were removed to a clean cage and received an injection of capsaicin (10 mg/kg, s.c. or i.p.). During the 5-minute post-injection period of social isolation, the arena was cleaned as above. Mice were then reintroduced to the same three-chamber apparatus and allowed to explore for an additional 15 minutes.

#### Experiment testing the effect of optogenetic activation of Agrp neurons on social investigation

For this experiment, we conducted the recordings on a modified testing chamber (25 × 35 × 23 cm) that allowed the quantification of social investigation in tethered mice. The test consisted of four phases: (1) habituation: test mouse was placed in the chamber with siblings for 10 minutes to acclimate to the optic cable; (2) baseline session: mouse was transferred to the central compartment and allowed to explore both end chambers for 15 minutes with the blue laser off; (3) opto-stimulation period: the mouse was returned to the sibling chamber and received 20 Hz stimulation (10 ms pulse width; 1 s on/1 s off) for 5 minutes via a 473 nm LED source (≤5 mW at the fiber tip); and (4) post-stimulation session: immediately following stimulation, the mouse was placed back in the central chamber for a final 15-minute exploration under continuous optogenetic stimulation (20 Hz, 10 ms pulses). Time in each chamber and investigation of the barrier were quantified as above.

### Statistical analysis and data plotting

The following software were used to analyze the data and plot the figures: mindthegraph, Biorender, Adobe Illustrator 2025, Prism 10, MATLAB 2024b, Microsoft Excel, and R. To compare two groups, we used *t* test. When analyzing three or more groups, we used a combination of one-way analysis of variance (ANOVA) or two-way ANOVA with repeated measures when data were paired. Holm-Sidak’s multiple comparisons test was then performed to find differences among groups. When necessary to calculate 95% confidence interval for each multiple comparison, Sidak’s test was used instead. The Geisser-Greenhouse correction for sphericity was used if necessary. To analyze the changes in response to specific stimuli related to the age of the animals, we used linear regression analysis to calculate the *r* square. Statistical data are provided in the figures and figure legends. A *P* value less than 0.05 was considered statistically significant.

