## Supplementary material for "Age-specific regulation of sociability by hypothalamic Agrp neurons": Figure S1

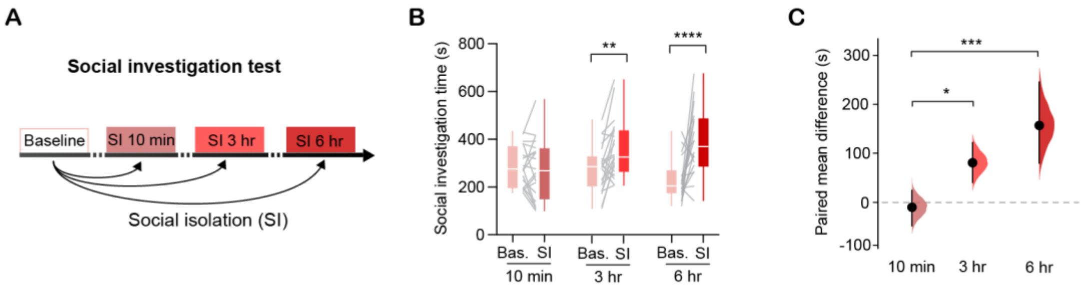


**Figure S1: Social isolation increases social investigation in juvenile mice.**

**(A)** Experimental design: juvenile mice (P24) were tested for baseline social investigation during an initial 15-minute test. Afterward, social investigation was re-assessed following periods of social isolation lasting either 10 minutes, 3 hours, or 6 hours. Stimulus mice were isolated for the same duration as the test animals before each social investigation test. **(B)** Effect of social isolation ($F_{1,\,57}$ = 21.57, $P < 10^{-5}$), social isolation duration ($F_{2,\,57}$ = 0.8, $P$ = 0.41) and interaction ($F_{2,\,57}$ = 8.45, $P$ = 0.0006). Number of animals: 10 min, n = 20; 3 h, n = 22; 6 h, n = 18). Statistical analysis using repeated measures two-way ANOVA followed by Holm-Sidak's multiple comparisons test. **(C)** Estimation plot displaying the paired mean difference (± 95% CI) of social investigation time after SI minus pre-SI at different isolation periods ($F_{2,\,57}$ = 8.48, P = 0.0006; one-way ANOVA followed by Holm-Sidak's multiple comparisons test). Box plot denotes minimum, first quartile, median, third quartile, and maximum values. Gray lines connecting box plots represent paired data. * $P$ < 0.05; ** $P$ < 0.01; *** $P$ < 0.001; **** $P$ < 0.0001.
