## Supplementary material for "Age-specific regulation of sociability by hypothalamic Agrp neurons": Figure S2

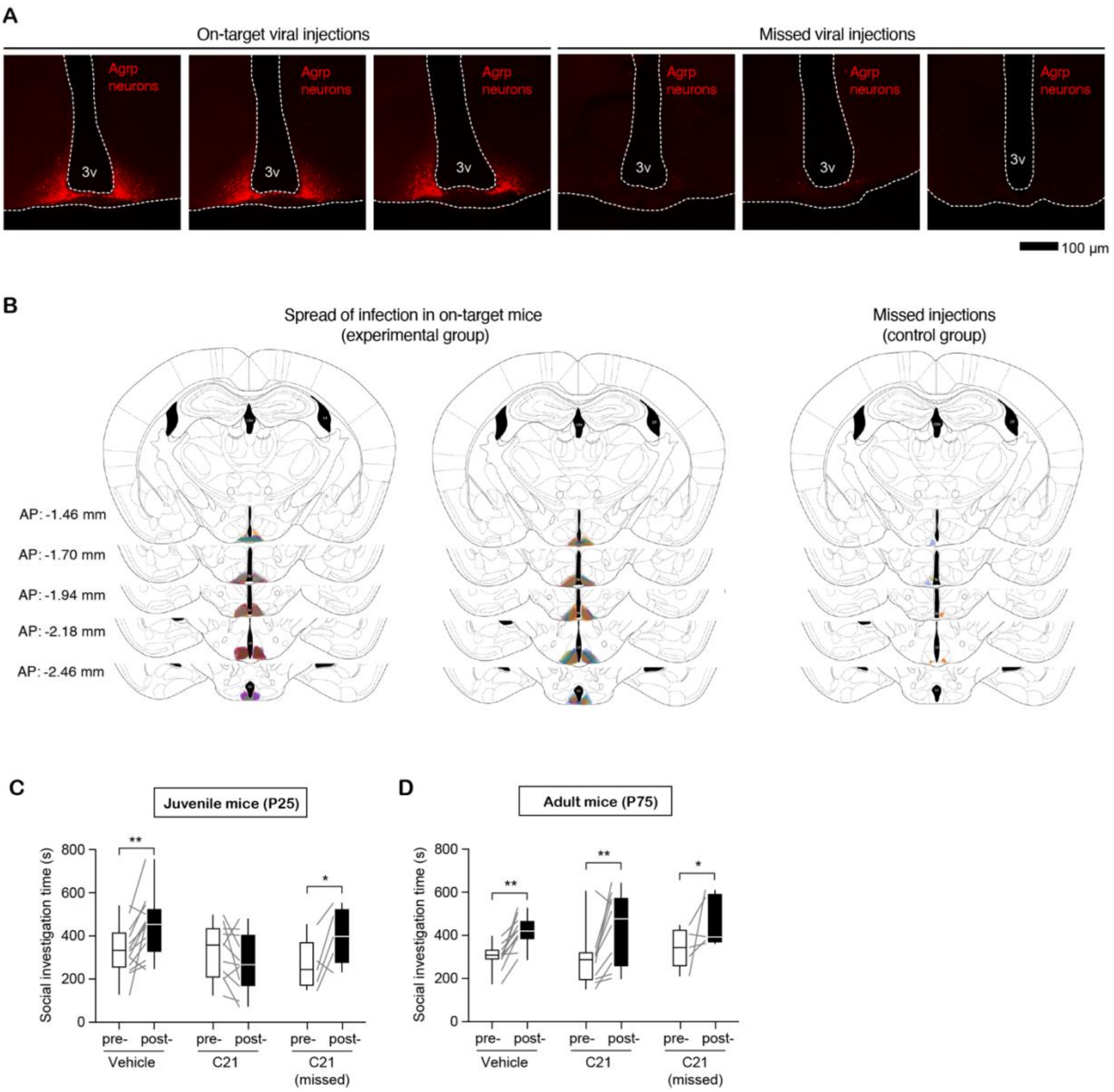


**Figure S2: Inhibition of Agrp neurons during social isolation blunts the increased sociability in juveniles, but not in adults. (A)** Representative images of mCherry labeling in Agrp neurons of mice injected with AAV-DIO-hM4Di-mCherry; three examples of mice showing good bilateral infection and three examples of mice showing missed injections. Mice with missed injections were used as controls (received C21 treatment). **(B)** Illustration of the expression of mCherry (hM4Di) in Agrp neurons of all mice used in this experiment. **(C)** Effect of social isolation for three hours ($F_{1, 26}$ = 8.11, $P$ = 0.008) and group ($F_{2, 26}$ = 1.83, $P$ = 0.17) on social investigation (interaction, $F_{2, 26}$ = 8.00, $P$ = 0.002) in juveniles (vehicle, n = 13; C21, n = 11, missed viral injections, C21, n = 5). **(D)** Similar to C, but in adults; effect of social isolation ($F_{1, 23}$ = 27.38, $P$ < 0.0001), group ($F_{2, 23}$ = 0.41, $P$ = 0.66) and interaction ($F_{2, 23}$ = 0.16, $P$ = 0.84) on social investigation (vehicle, n = 11; C21, n = 10, missed viral injections, C21, n = 5). In C and D: statistical analyses using repeated measures two-way ANOVA followed by Holm-Sidak's multiple comparisons test. Box plot denotes minimum, first quartile, median, third quartile, and maximum values. Gray lines connecting box plots represent paired data. * $P$ < 0.05; ** $P$ < 0.01; *** $P$ < 0.001; **** $P$ < 0.0001.
