## Supplementary material for "Age-specific regulation of sociability by hypothalamic Agrp neurons": Figure S3

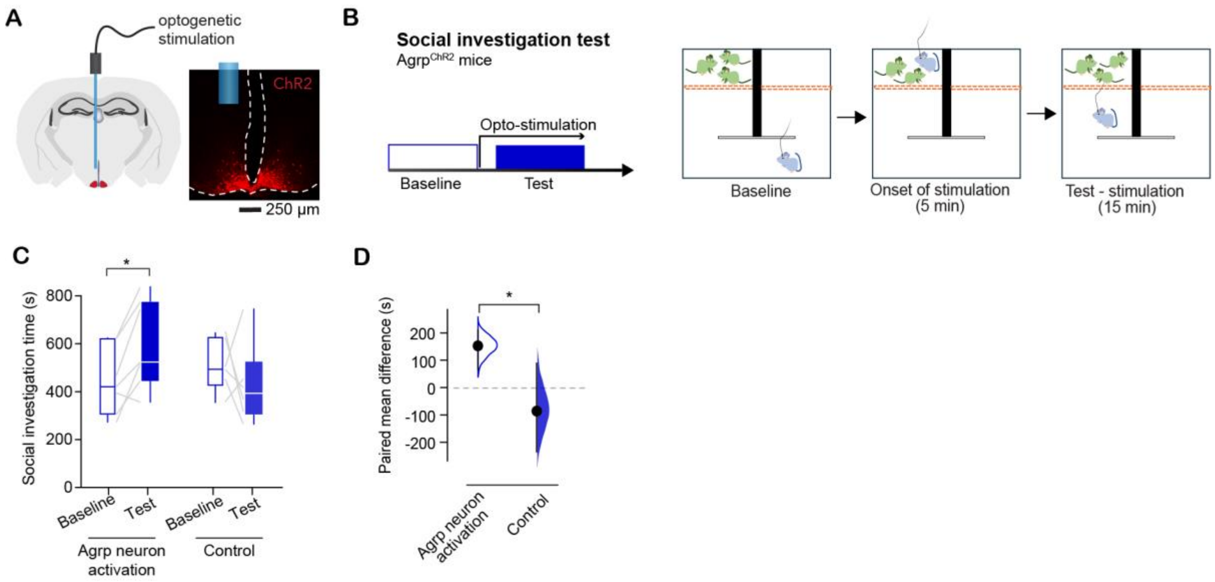


**Figure S3: Optogenetic stimulation of Agrp neurons increases social investigation in juvenile mice. (A)** Representative image of fiber optic cannula placement in $Agrp^{ChR2}$ mice and expression of ChR2-mCherry in Agrp neurons. **(B)** Experimental design to test the effect of optogenetic stimulation of Agrp neurons in a test of social investigation in juvenile mice. First, the experimental mouse was allowed to explore the recording chamber for 15 minutes; the U-shaped chamber contained two compartments, with a mesh barrier at the end—at one end there was a group of cage mate siblings. After this initial baseline period, the experimental mouse was returned to the sibling chamber and received 20 Hz opto-stimulation (10 ms pulse width; 1 s on/1 s off) for 5 minutes via a 473 nm LED source (≤5 mW at the fiber tip). Immediately following stimulation, the mouse was placed back in the central chamber for a final 15-minute exploration under optogenetic stimulation (20 Hz, 10 ms pulses, as before). **(C)** There was no significant overall main effect of opto-stimulation ($F_{1, 11}$ = 0.52, $P$ = 0.48) or group ($F_{1, 11}$ = 0.47, $P$ = 0.50), but there was significant effect of their interaction ($F_{1, 11}$ = 6.77, $P$ = 0.02), indicating that optogenetic stimulation of Agrp neurons differentially affected social investigation during the test phase. Number of mice: $Agrp^{ChR2}$, n = 7; control, n = 6. Statistical analysis using repeated measures two-way ANOVA followed by Fisher's multiple comparisons test. **(D)** Estimation plot displaying the paired mean difference (± 95% CI) of social investigation time comparing the period of opto-stimulation minus the baseline period in control and Agrp neuron activated mice ($t_{11}$ = 2.6, $P$ = 0.02; unpaired *t*-test). In C: box plot denotes minimum, first quartile, median, third quartile, and maximum values. Gray lines connecting box plots represent paired data. * $P$ < 0.05; ** $P$ < 0.01; *** $P$ < 0.001; **** $P$ < 0.0001.
