## Supplementary material for "Age-specific regulation of sociability by hypothalamic Agrp neurons": Figure S4

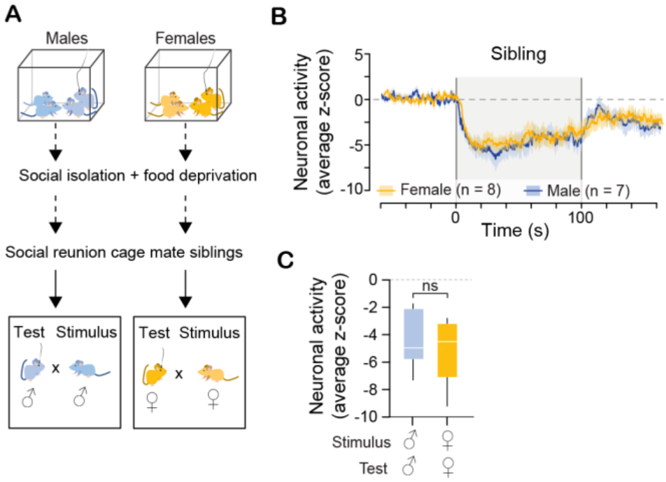


**Figure S4: The sex of the interacting sibling does not affect the response of Agrp neurons to social reunion in juvenile mice. (A)** Juvenile mice (P22-P28) were socially isolated, and food deprived for 3-5 hours, then allowed to interact with their same sex cage mate sibling for 100 seconds. **(B)** The average Z score of Agrp neuron activity; shaded area represents the period during which a same sex cage mate sibling was present in the recording chamber. Lines represent mean ± sem. **(C)** Comparison of the mean response of male and female mice to same sex cage mate siblings ($t_{13}$ = 0.57, $P$ > 0.05; unpaired $t$-test). Box plot denotes minimum, first quartile, median, third quartile, and maximum values.
