## Supplementary material for "Age-specific regulation of sociability by hypothalamic Agrp neurons": Figure S5

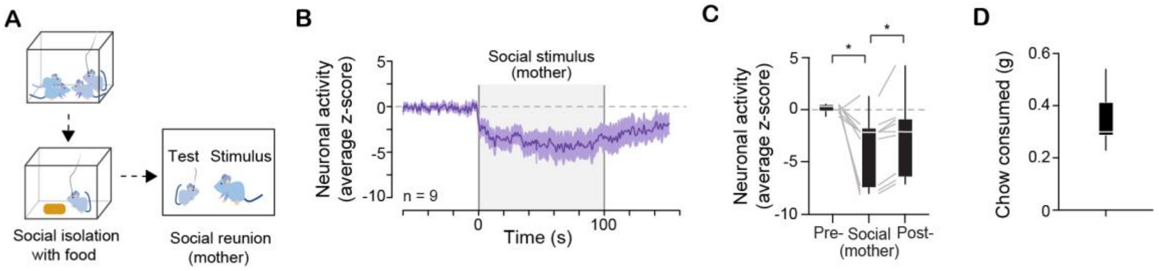


**Figure S5: Social reunion reduces Agrp neuron activity in juvenile mice that are socially isolated, but not food deprived. (A)** Juvenile mice (P22-P28) were socially isolated with ad libitum food for 3-5 hours, then allowed to interact with their mother as a social stimulus for 100 seconds. **(B)** The average Z score of Agrp neuron activity; shaded area represents the period during which the mother was present in the recording chamber (n = 9). Lines represent mean ± sem. **(C)** The mean Z score during baseline, social interaction, and recovery (ε = 0.53, $F_{1.06,\ 8.50} = 7.22$, $P = 0.02$; repeated measures one-way ANOVA with Geisser-Greenhouse correction followed by Holm-Sidak's multiple comparisons test). **(D)** Amount of food consumed during isolation period. In C and D: box plot denotes minimum, first quartile, median, third quartile, and maximum values. Gray lines connecting box plots represent paired data. * $P < 0.05$.
