## Supplementary material for "Age-specific regulation of sociability by hypothalamic Agrp neurons": Figure S6

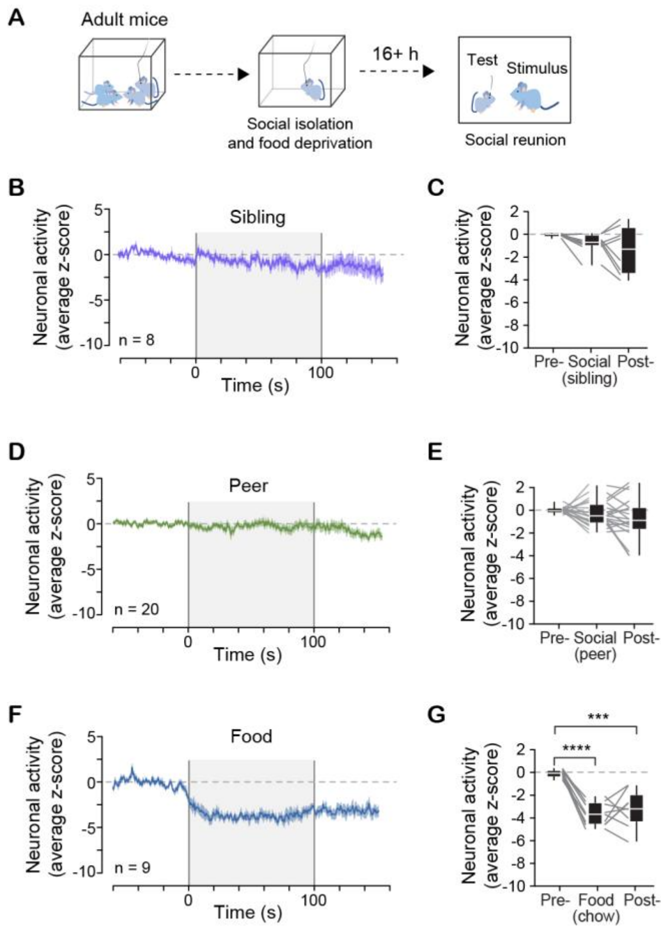


**Figure S6: Adult mice do not respond to social reunion even after a prolonged period of social isolation and food deprivation. (A)** Adult mice were socially isolated, and food deprived overnight for 16 hours, then allowed to interact with their same sex cage mate siblings and peers; subsequently they were presented with food as control. All stimuli were presented for 100 seconds. **(B)** The average Z score of Agrp neuron activity; shaded area represents the period during which a sibling was present in the recording chamber. **(C)** Related to B, the mean Z score during baseline, social interaction, and recovery ($\varepsilon = 0.53$, $F_{1.18,\ 8.32} = 3.03$, $P = 0.11$). **(D)** The average Z score of Agrp neuron activity; shaded area represents the period during which a peer was present in the recording chamber. **(E)** Related to D, the mean Z score during baseline, social interaction, and recovery ($\varepsilon = 0.70$, $F_{1.14,\ 26.85} = 3.005$, $P = 0.08$). **(F)** The average Z score of Agrp neuron activity; shaded area represents the period during which food was present in the recording chamber. **(G)** Related to F, the mean Z score during baseline, food presentation, and recovery ($\varepsilon = 0.72$, $F_{1.44,\ 11.57} = 35.99$, $P = 0.00002$). In B, D, and F: lines represent mean ± sem. In C, E, and G: repeated measures one-way ANOVA with Geisser-Greenhouse correction followed by Holm-Sidak's multiple comparisons test. Box plot denotes minimum, first quartile, median, third quartile, and maximum values. Gray lines connecting box plots represent paired data. * $P < 0.05$; ** $P < 0.01$; *** $P < 0.001$; **** $P < 0.0001$.
